# Molecular analysis of the midbrain dopaminergic niche during neurogenesis

**DOI:** 10.1101/155846

**Authors:** Enrique M. Toledo, Gioele La Manno, Pia Rivetti di Val Cervo, Daniel Gyllborg, Saiful Islam, Carlos Villaescusa, Sten Linnarsson, Ernest Arenas

## Abstract

Midbrain dopaminergic (mDA) neurons degenerate in Parkinson’s disease and are one of the main targets for cell replacement therapies. However, a comprehensive view of the signals and cell types contributing to mDA neurogenesis is not yet available. By analyzing the transcriptome of the mouse ventral midbrain at a tissue and single-cell level during mDA neurogenesis we found that three recently identified radial glia types 1-3 (Rgl1-3) contribute to different key aspects of mDA neurogenesis. While Rgl3 expressed most extracellular matrix components and multiple ligands for various pathways controlling mDA neuron development, such as Wnt and Shh, Rgl1-2 expressed most receptors. Moreover, we found that specific transcription factor networks explain the transcriptome and suggest a function for each individual radial glia. A network controlling neurogenesis was found in Rgl1, progenitor maintenance in Rgl2 and the secretion of factors forming the mDA niche by Rgl3. Our results thus uncover a broad repertoire of developmental signals expressed by each midbrain cell type during mDA neurogenesis. Cells identified for their emerging importance are Rgl3, a niche cell type, and Rgl1, a neurogenic progenitor that expresses *ARNTL,* a transcription factor that we find is required for mDA neurogenesis.

## INTRODUCTION

Our knowledge of the contribution of individual signals to midbrain dopaminergic (mDA) neuron development has grown in a considerable manner in recent years. It is currently thought that the development of mDA neurons is controlled by the interaction of multiple transcription factors and signaling pathways (Arenas *et al*, 2015; Smidt & Burbach, 2007). However, we still lack the systematic knowledge required to understand how all these factors are coordinated over time and space, in the complex signaling microenvironments in which the cells reside. These microenvironments, referred to as niches, consist of an extracellular matrix (ECM) and local paracrine/autocrine signals contributed by neighboring cells (Jones & Wagers, 2008). Far from static, such microenvironments are constantly being remodeled (Scadden, 2006; Jones & Wagers, 2008), leading to progressive changes in composition over time, which control fundamental developmental processes such as cell fate or the balance between proliferation and neurogenesis (Lu *et al*, 2011). In the nervous system, some of the critical cell types controlling neural development are radial glia cells. Typically, radial glia (Rgl) cells are bipolar cells with their somas located in the ventricular zone (VZ) and their processes extending to the ventricular cavity and radially to the pial surface. These cells, once thought to be a scaffold for neurogenesis, are currently thought to be transient stem cells, capable of limited self-renewal, of giving rise to different cell lineages and of undergoing neurogenesis by asymmetrical cell division (Götz & Huttner, 2005; Taverna *et al*, 2014). In the developing VM, it is unknown how many cells contribute to the DA neurogenic niche, how they contribute to neurogenesis, what factors do they express and what are the transcriptional networks operating in each of these cells. We previously reported that a proliferative floor plate Rgl cell gives rise to mDA neurons through the generation of a lineage with subsequent cellular intermediates (Bonilla *et al*, 2008). However, we recently performed single-cell RNA-sequencing (RNA-seq) of the developing midbrain and identified multiple cell types, including three different types of molecularly-diverse radial glia (Rgl1-3) with distinct temporal gene expression patterns and spatial distribution (La Manno *et al*, 2016). This finding has thus raised several important questions such as: Which of these three radial glia cell types are capable of undergoing neurogenesis in the developing VM? Do they also contribute to neurogenesis with cell extrinsic factors? How are cell intrinsic and extrinsic factors spatially and temporally integrated? What is the cellular microenvironment or niche in which mDA neurogenesis takes place?

In order to address these questions, we used single cell and bulk RNA-sequencing as well as systems biology methodologies to gain further understanding of the cellular and molecular components of the mDA neurogenic niche. Our study identifies Rgl1-3 as the main cell types contributing signals and cells to mDA neurogenesis. However, each cell type contributed in a different way. While Rgl1 was identified as the neurogenic radial glia cell type, Rgl3 was the main contributor to cell extrinsic factors in the mDA neurogenic niche, including morphogens, growth factors and ECM. Our work thus provides the first unbiased and integrated analysis of the major molecular processes taking place in the mDA neurogenic niche. We also found how the expression of transcription factors, as well as signaling and ECM components, are controlled over time and in defined cell types forming the mDA neurogenic niche. We conclude that efforts aiming at recapitulating mDA neuron development in stem cells, either for cell replacement or for in vitro disease modeling, should thus focus on the generation of Rgl1 and Rgl3, the two main pillars of mDA neurogenesis.

## RESULTS

### Transcriptomic analysis of the embryonic ventral midbrain

In order to obtain a genome-wide view of the niche occupied by midbrain progenitor cells during mDA neurogenesis, we performed bulk RNA-seq of the mouse VM and surrounding tissues from embryonic day (E)11.5 to E14.5, the period at which mDA neurogenesis takes place (Arenas *et al*, 2015). We obtained transcriptomic profiles from five different regions of the neural tube of TH-GFP mouse embryos (Matsushita *et al*, 2002): the VM, the ventral hindbrain (HB), ventral forebrain (FB), dorsal midbrain (DM) and alar plate (L) (Fig. 1A). Specific anatomical and temporal gene expression patterns were identified in all samples (Fig. 1B). A pair-wise correlation comparing all the VM samples revealed that E11.5 was the most divergent stage (Fig. 1C). Furthermore, principal component analysis (PCA) confirmed the similarity between E12.5 and E13.5 (Fig. 1D, Fig. EV1A). Multiple differentially expressed genes (DEG) were identified in the VM, compared to the other dissected brain regions (Table EV1). The VM transcriptome was further analyzed by weighted gene co-expression network analysis. Eight gene modules were identified and found to correlate with distinct expression patterns that describe the changes in transcriptome over time (Fig. EV1B-D, details in Table EV2). These modules can be summarized in three general developmental patterns. The first pattern (Fig. 1E, three modules in EV1B), consists of genes whose expression is increased over time, such as *Slc6a3* (DAT, dopamine transporter), a marker of mDA neuron maturation (Fauchey *et al*, 2000). Gene ontology (GO) analysis of this module showed enrichment in processes related to neuronal development and maturation, peaking at E14.5 (Fig. EV1F). The second pattern consists of genes whose expression peaks at middle time points E12.5- 13.5 (Fig. 1F, two modules in EV1D), such as *Ncor2*, which represses the differentiation of neuronal precursors (Jepsen *et al*, 2007). GO analysis showed enrichment of genes involved in tissue growth and developmental processes (Fig. EV1E). Lastly, we identified a third pattern with decreasing gene expression during development, from E11.5 (Fig. 1G, three modules in EV1C). This included genes such as *Notch3* and *Hes5*, a ligand and an effector of the Notch signaling pathway, respectively (Louvi & Artavanis-Tsakonas, 2006). GO analysis showed enrichment in processes that describe cell proliferation (Fig. EV1G). These analyses thus provided both a detailed molecular insight and a general overview of the biological processes taking place in the developing VM during mDA neurogenesis, in which there is a shift from proliferation to progressive neuronal maturation and repression of progenitor pathways and functions.

**Figure 1.**
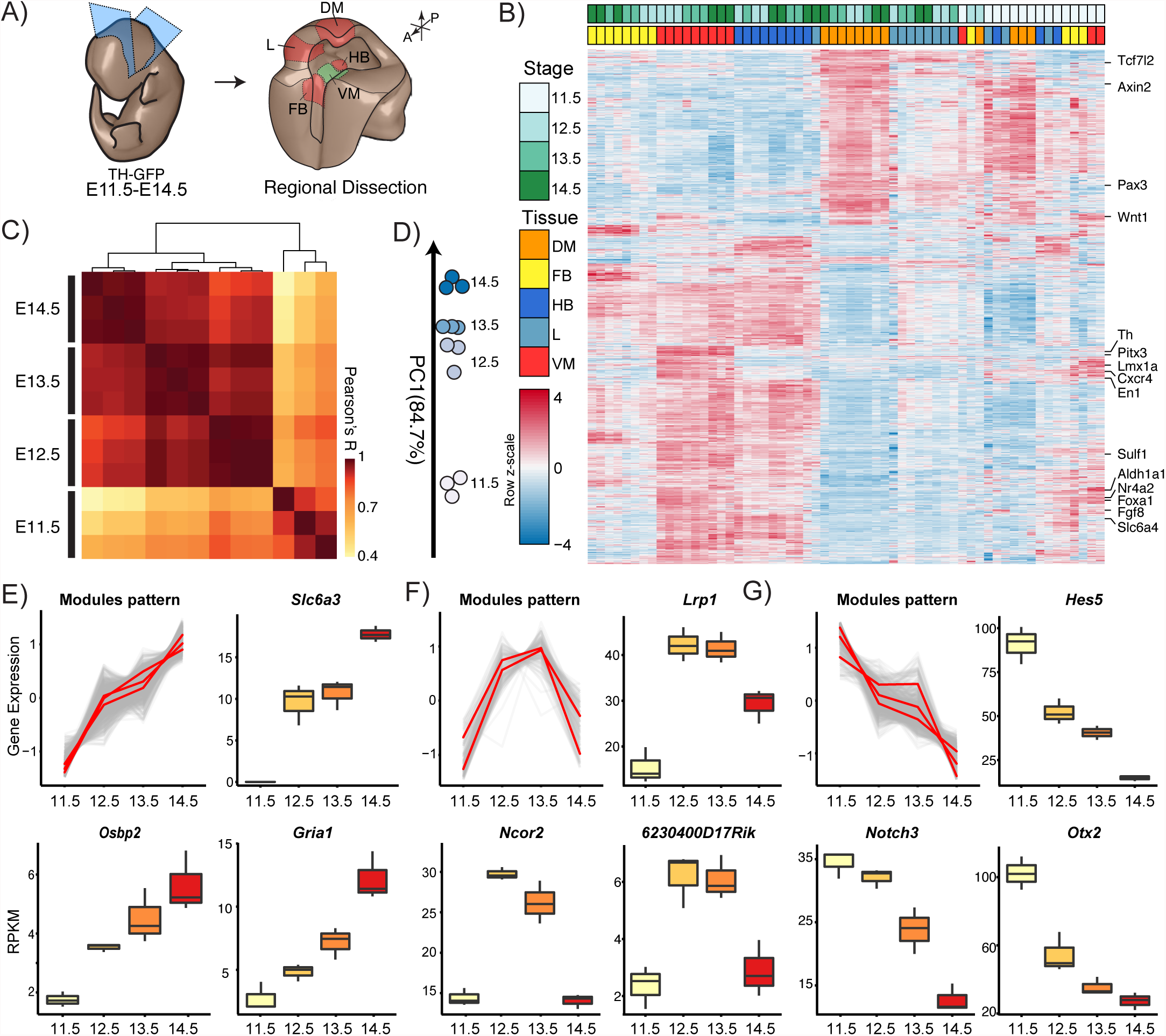
Transcriptomic profile of the ventral midbrain. **(A)** Embryonic tissue dissection scheme and regions collected for RNA-seq from TH-GFP embryos of stages E11.5, E12.5, E13.5 and E14.5. VM, ventral midbrain. DM, dorsal midbrain. FB, forebrain floor plate. HB, hindbrain floor plate. L, alar plate. **(B)** Heat map representation of high variance genes (rows) in dataset of all samples (columns) across all time points. Samples are color coded for stage and region. **(C)** Correlation of VM samples after filtering for variance (12.5%). **(D)** First principal component analysis of VM samples across all time points. Full plot is in Fig. EV1A. (E-G) Gene expression patterns in the developing VM as obtained by WGCNA of the VM samples. **(E)** Pattern summarizing gene modules with positive correlation over development (top left) and examples of genes: *Slc6a3*, *Ostbp2* and *Gria1*. **(F)** Pattern summarizing gene modules correlating at E12.5 and E13.5 and examples of genes in these modules: *Lrp1*, *Ncor2* and *6230400d17Rik*. **(G)** Pattern summarizing gene modules with negative correlation over development and some examples: *Hes5*, *Notch3* and *Otx2*. Expression is in RPKM. Red lines represent the median expression of each module. All modules with significant correlations are shown in Fig. EV1B-D.

### The dopaminergic module: A network that defines the midbrain dopaminergic neuron niche

To identify molecular processes that specifically occur during VM and mDA neurons, we performed a second unbiased weighted gene co-expression network analysis of the transcriptomes, but this time including both the VM and neighboring regions. This analysis generated 13 modules (or networks) of co-expressed genes, one being the “light green” module (Fig. EV2A) that had the most enriched DEG in the VM from E11.5 to E14.5 (Fig. EV2B and Table 1). Further analysis revealed that the top 5% of the interactions (1235 out of 24506) involved 181 genes (out of 374), which were enough to separate the VM samples at all the stages analyzed, as assessed by PCA (Fig. EV2C). This refined module contained many known genes expressed in the mDA lineage (*Foxa1*, *Shh*, *Wnt5a*, *Th*, *Ddc*, *Snca*, etc) and was thus designated as the mDA module. This module was represented as an undirected co-expression network (Fig. 2A), in which the color of the node indicates changes in gene expression (RPKM) over time and the lines linking them represent the pair-wise correlation of the linked genes (Langfelder & Horvath, 2008). GO analysis of the mDA module provided us with an overview of the biological processes in which these genes participate. These included relatively well-studied processes in mDA neuron development, such as dopamine biosynthetic process (p-value: 4.19e-9), morphogenesis of epithelium (p-value: 2.85e-5) or neural tube development (p-value: 1.91e-3); but also less well studied processes such extracellular matrix (p-value: 2.17e-7) or extracellular matrix component (p-value: 8.04e-3), amongst others (Fig. EV2D). These results indicate that the ECM is likely to play a more significant role than previously anticipated in mDA neuron development.

**Figure 2.**
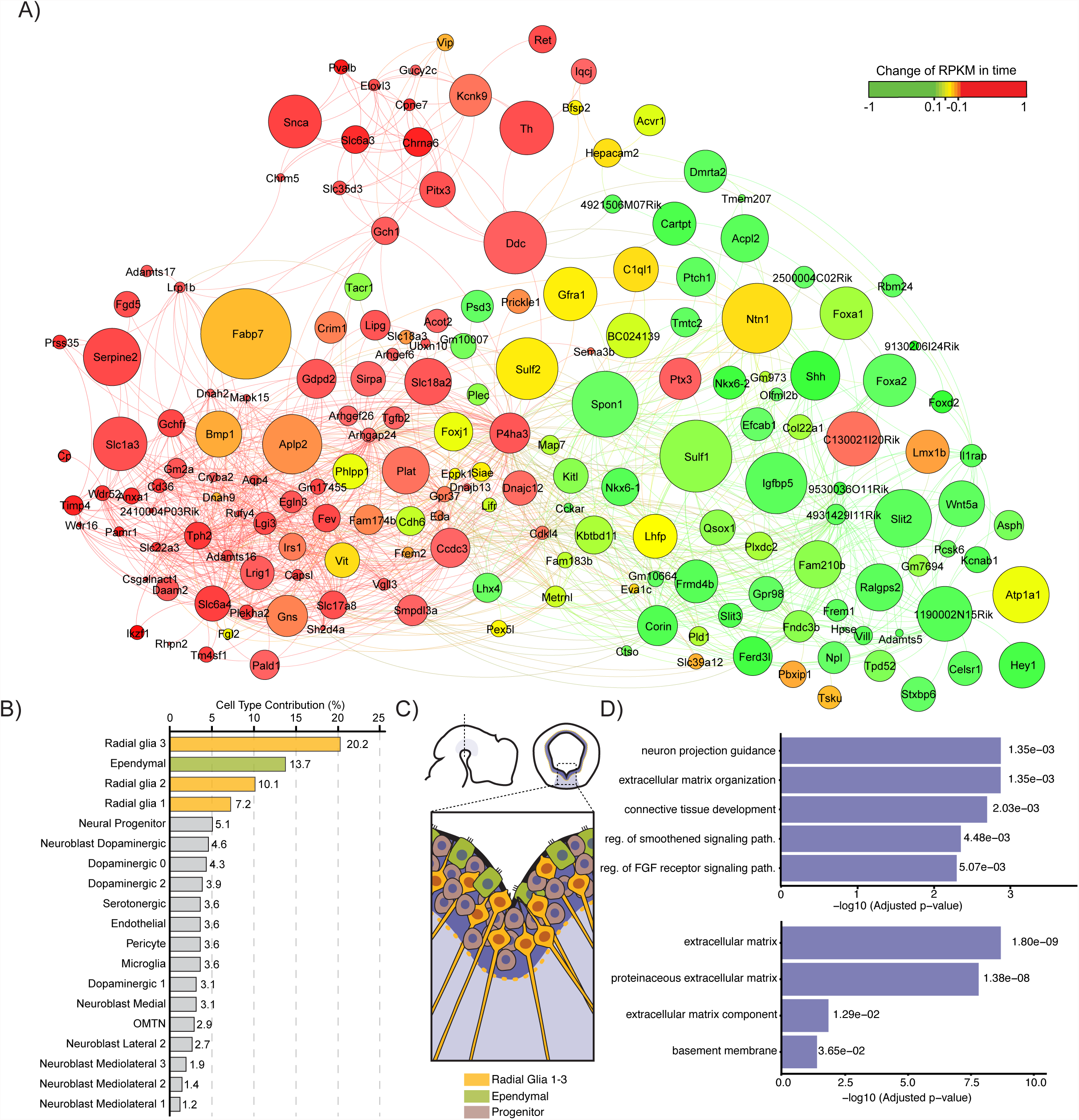
The dopaminergic module and its deconvolution at a single-cell level. **(A)** Weighted gene co-expression network analysis of the mDA module, filtered for the top 5% of interactions. Color represents changes in levels of gene expression during development. Node size is proportional to the mean expression levels of the gene during development. **(B)** Contribution of the different cell types in the VM to the network formed by the dopaminergic module. **(C)** Diagram of VM floor plate with focus on the cell types present in the VZ**. (D)** GO enrichment terms corresponding to genes with the highest contribution to the mDA module. Top, GO for biological process. Bottom, GO for cellular components.

**Table 1.**

| Module | Number VM<br>DEG | Module size | q-value | Enrichment<br>Score |
| --- | --- | --- | --- | --- |
| Black | 37 | 551 | 9.12e-02 | -0.55 |
| Brown | 46 | 856 | 1.08e-01 | -0.14 |
| Cyan | 6 | 281 | 9.15e-04 | -5.68 |
| Green | 119 | 2839 | 9.26e-03 | 6.10 |
| Green Yellow | 10 | 423 | 1.80e-04 | -6.35 |
| Grey | 24 | 519 | 3.99e-02 | -1.53 |
| Grey60 | 55 | 175 | 1.03e-24 | 5.80 |
| Light cyan | 18 | 201 | 3.26e-02 | -2.01 |
| Light green | 191 | 378 | 5.88e-136 | 824.97 |
| Light yellow | 2 | 100 | 2.46e-02 | -3.28 |
| Magenta | 69 | 3087 | 3.56e-17 | 13.90 |
| Salmon | 54 | 287 | 6.10e-14 | 2.63 |
| Tan | 11 | 371 | 2.63e-03 | -4.27 |

In order to identify the cell types contributing to the mDA module by gene expression, we used a single-cell RNA-seq analysis of the VM that recently allowed us to identify the twenty-six cell types present in the mouse VM during mDA neurogenesis (La Manno *et al*, 2016). This dataset was used to perform a cell type deconvolution of our mDA network, assigning cell type/s to the identified expression profiles in the mDA network (Fig. EV3A). 97% of the genes in the network (176 of 181 genes) were assigned to at least one cell type. Most of the genes (51.2%) were contributed by only four cell types present in the VZ: three types of radial glia like cells (Rgl 1-3) and ependymal cells (Fig. 2B, C). Sixteen additional cell types expressed the remaining 48.8% of the gene on the mDA network, while the remaining six other cell types did not contribute to this network. Notably, the four VZ cell types were the main contributors to the formation of the ECM (p-value: 7.72e-26) and its regulation (p-value: 2.88e-15), according to the gene sets defined in the Matrisome project (http://matrisomeproject.mit.edu/) (Naba *et al*, 2012). GO enrichment analysis confirmed that the main contribution of the four VZ cells types was in genes of ECM components, neural projections as well as in Shh and FGF signaling pathways (Fig. 2D). These results thus identify Rgl and ependymal cells as important contributors to the expression of ECM and critical signaling factors controlling mDA neuron development.

We next examined the specific contribution of the most abundant cell types in the mDA niche during neurogenesis, that is, radial glia, but not ependymal cells, which emerge later. Radial glia has been previously identified as a neurogenic cell in the mDA lineage (Bonilla *et al*, 2008) and as the source of multiple signaling molecules in the midbrain floor plate (Arenas *et al*, 2015). We thus decided to compare the transcriptome profiles of individual radial glia cell types, Rgl1-3, by gene set enrichment analysis (Subramanian *et al*, 2005) in order to explore whether distinct radial glia cell types may be poised to serve distinct functions. Analysis of Rgl1 revealed an enrichment in the expression of genes related to cell proliferation, such as M Phase (Normalized Enrichment Score (NES): 2.341, q-value <0.001), ribosomal constituents (NES: 2.641, q-value <0.001) and cellular biosynthetic process (NES: 1.992, q-value <0.001), which defined Rgl1 as a cell in a highly proliferative state. Rgl2, also expressed genes related to proliferation (Mitosis, NES: 1.759, q-value: 0.04), but additionally expressed genes involved in fatty acid metabolism (NES 1.837, q-value: 0.01) and cholesterol biosynthesis (NES: 1.893, q-value: 0.006), possibly linking Rgl2 to the production of specific midbrain cholesterol metabolites controlling mDA neurogenesis (Theofilopoulos *et al*, 2013). The last radial glia, Rgl3, was not enriched for proliferation genes, but rather in ECM core components (NES 2.029, q-value <0.001) and ECM regulators (NES 2.363, q-value <0.001) as well as signaling pathways (Glypican 1 Pathway, NES: 1.840, q-value: 0.02, Netrin 1 signaling, NES: 1.809, q-value: 0.04). These results suggested a high degree of specialization of radial glia and lead us to examine in further detail their contribution to the mDA niche with regards to secreted signaling molecules, such as morphogens and growth factors as well as ECM.

### The extracellular matrix in the ventral midbrain

In order to further delve into the possible role of the ECM, we decided to identify the VM cell types responsible for the expression of genes coding for critical components of the ECM, regardless of their contribution to the mDA module. In order to rank the cell type contribution to the ECM, we developed a score based on the number of genes being expressed and their levels of expression (see methods), taking in account both core ECM components and ECM regulators, as defined by the Matrisome gene sets (Hynes & Naba, 2012; Naba *et al*, 2012). Six cell types (Rgl2-3, ependymal cells, endothelial, pericytes and microglia) were above a threshold set at 99.9% quantile of the mean ECM score (Fig. 3A). These cell types contributed to 82.8% of the total number of mRNA molecules of ECM core components (Fig. EV4A), and to 88.2% of ECM regulators in the VM (Fig. EV4B). Genes coding for core ECM components (Fig. 3B, EV4C) or ECM modifiers (Fig. 3C, EV4D) that were significantly expressed above baseline are shown in a color scale proportional to mRNA molecules for each cell type. Our results show that Rgl3, endothelial cells and pericytes were the main contributors to ECM core transcripts in the VM (Fig. 3A). While these 3 cell types shared some of the transcripts (*Sparc* and *Sparccl1*), each cell type expressed specific transcripts, such as *vitronectin, colagen4a1/2* and *agrin* in endothelia and pericytes or *netrin1, decorin* and *spondin1* in Rgl3 (Fig. 3B). The latter finding is particularly interesting as we identified Rgl3 as a key component of the mDA neurogenic niche (Fig. 2) and its products, Netrin1 and Decorin are known to regulate axonal development in mDA neurons (Kastenhuber *et al*, 2009) and midbrain neurogenesis (Long *et al*, 2016), respectively. On the other hand, we found that microglia is the main cell type involved in ECM regulation (Fig. 3D), in particular, by expressing multiple types of cathepsin proteases (Fig. 3C), which likely contribute to remodel the ECM and allow them to migrate through the brain parenchyma (Chapman *et al*, 1997). Finally, from a temporal perspective, we found that Rgl1 and pericytes are more abundant during early neurogenesis (E11.5-12.5) while Rgl2-3 predominates during later stage (E14.5-E15.5) and ependymal cells subsequently after that at E18.5 (Fig. 3E). Adjusting the scores to the relative abundance of cell types at every stage confirmed the importance of the six cell types above and indicated an additional contribution of immature DA neurons (DA0) and Neuronal progenitors (NProg) at E11.5-E12.5 (Fig. S5 A-F). In summary, our results suggest a quantitatively important and constant contribution to the ECM by non midbrain-specific cell types, such as endothelial cells pericytes and microglia, suggesting a generic role of these cells. Instead, we found a midbrain- and stage-specific early contribution of NProg and Rgl1, and a late one by Rgl3.

**Figure 3.**
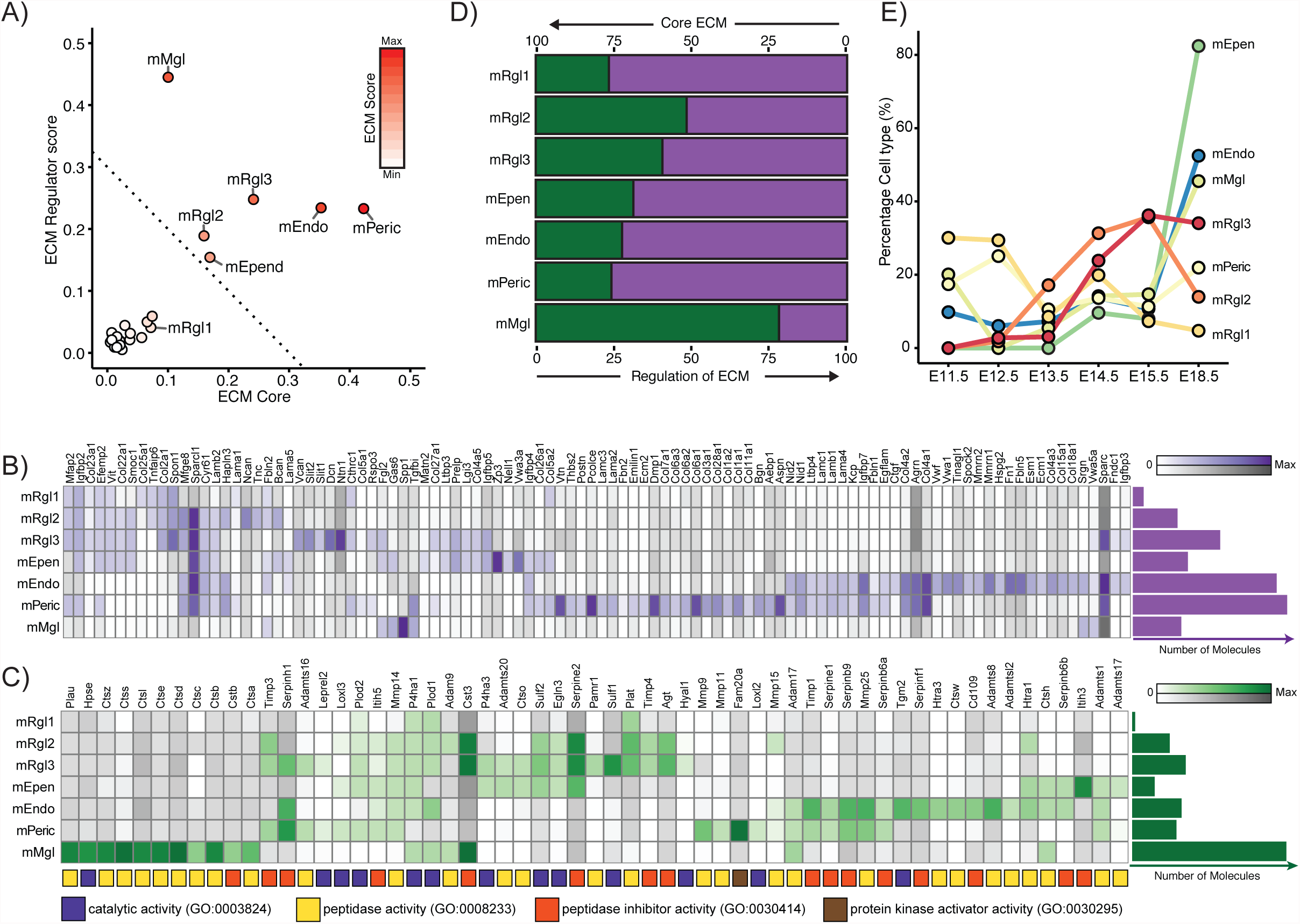
Analysis of the contribution of individual mouse VM cell types to the composition and dynamics of the extracellular matrix. **(A)** Cell type contribution to the ECM as determined by scores for ECM core components and ECM regulators. Dotted line represents quantile 99.9% of the bootstrapped mean of the ECM scores. **(B)** Heat map of the expression of core components of the ECM by the most significant cell types. Color intensity is proportional to the Bayesian estimate of expression level. Gray scale indicates values below significance level. mRgl1-3, radial glia type 1 to 3; mPeri, pericyte; mEndo, endothelial cell; mEpend, ependymal cell; mMgl, microglia. **(C)** Heat map of the expression of genes regulating the ECM regulator genes by the most significant cell types, as described as above. Color boxes under each gene identify their ontology. (**B, C**) The horizontal plots represent the total number of molecules per cell type. **(D)** Percentage of genes expressed by each cell type that participate in the regulation of the ECM (green) versus core ECM components (purple). **(E)** Relative abundance of cell types at different developmental stages, as sampled by single cell RNA-sequencing of the VM (La Manno *et al*, 2016).

### Signaling pathways in the ventral midbrain

Our analysis of the mDA module in addition of suggesting an important contribution of radial glia to the ECM, also identified the cell types expressing signaling pathway components known to participate in mDA neuron development (Fig. 2B-D). To determine the possible directionality of the developmental signaling pathways, we curated a gene set of ligands and their corresponding receptors (see methods, Table EV3) for known signaling pathways and used that gene set to identify the cell types expressing ligands and their receptors in the VM. As in the case of the ECM, we examined the involvement of a cell type in signaling calculating a receptor score and a ligand score (see methods). This analysis identified seven cell types (Rgl1-3, ependymal cells, endothelial, pericytes and microglia) as the main contributors to signaling (Fig. 4A), covering multiple ligands (Fig. 4B) and receptor families (Fig. 4C), several of which are known for their role in embryonic midbrain development (Arenas *et al*, 2015; Smidt & Burbach, 2007). Notably, Rgl3 and pericytes showed the highest score for combined number and levels of ligands and receptors (Fig. 4A), which indicates that Rgl3 and pericytes are the main signaling centers in the VM. These cells offer the possibility of signaling in an autocrine/paracrine manner and of a cross-talk between the neural tissue and the vasculature, whose function and significance for mDA neuron development remains to be determined. On the other hand, Rgl1-2 expressed more receptors than ligands (Fig. 4A), which make them the main candidates to respond to and execute different aspects of VM and mDA neuron development.

**Figure 4.**
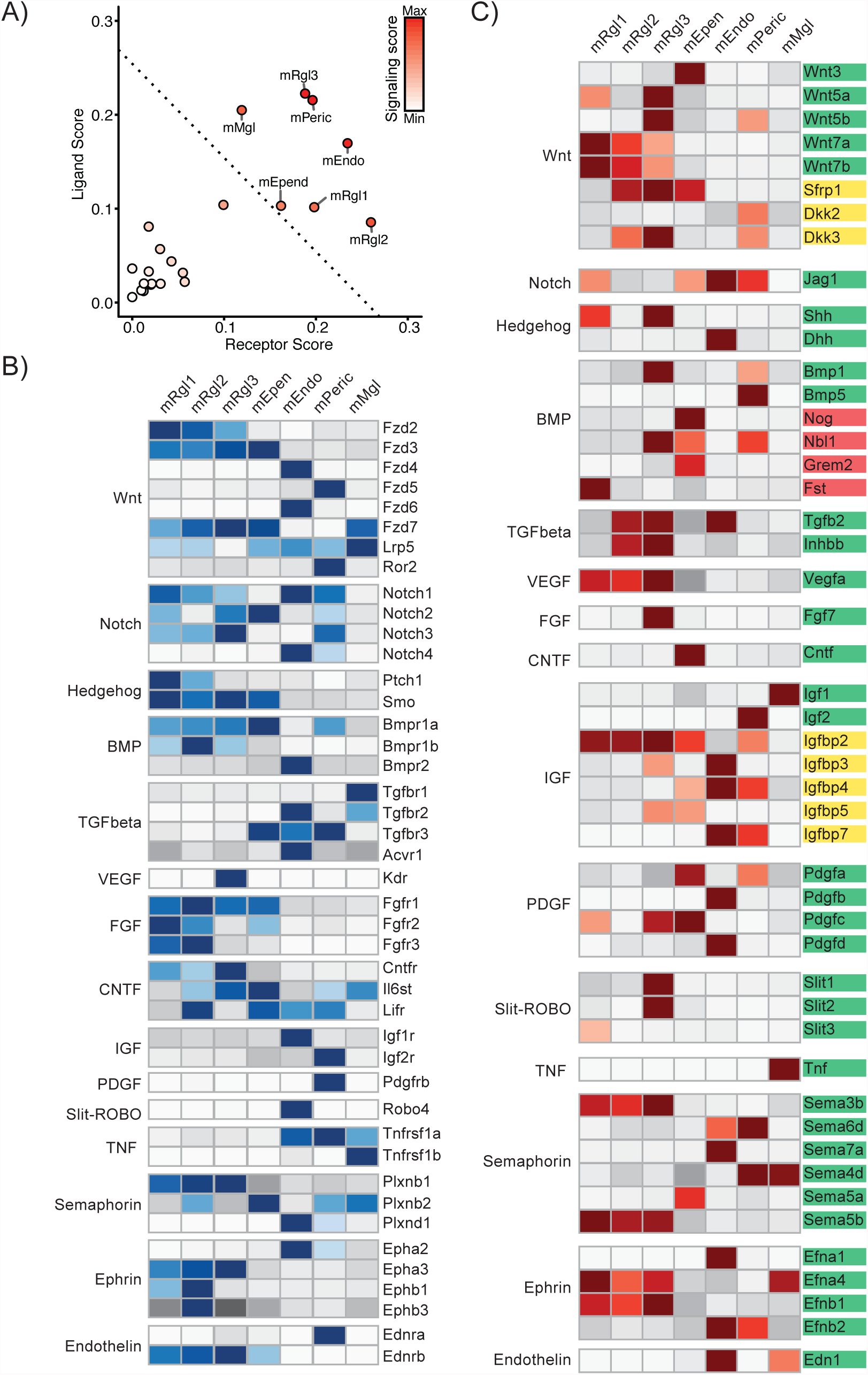
Contribution of individual mouse VM cell types to signaling in the mDA niche. **(A)** Plot showing the receptor and ligand scores of the different VM cell types. Color is proportional to the signaling score. Dotted line represents quantile 99.9% of the bootstrapped mean of the signaling scores. **(B)** Heat map representation of the expression of receptors in identified VM cell types. **(C)** Heat map representation of the ligands expressed by the most cell types in the VM. Gene colored according to their activity; green for activation, red for inhibition and yellow for context-dependent modulation. (B, C) Color intensity is proportional to the Bayesian estimate of expression level. Gray scale indicates values below significance level. mRgl1-3, radial glia type 1 to 3; mPeri, pericyte; mEndo, endothelial cell; mEpend, ependymal cell; mMgl, microglia.

As with the ECM score, we adjusted the signaling score to cell type abundance. This analysis suggests an early role of mRgl1 and progenitor cells at early stages (Fig. S5 G-I), and a late role of generic cell types such as endothelial cells and pericytes, as well as midbrain-specific cell types, such as mRgl2 and mRgl3 (Fig. S5 J-L).

Wnt signaling was represented with the highest number of types of ligands and receptors (Fig. 4B-C), underlining the prevalence of this pathway and its importance in mDA neuron development (Arenas, 2014). Interestingly, our analysis predicts novel candidate cell-to-cell signaling events with single-cell resolution (Fig. 4B-C). For instance, we found evidence for autocrine loops in multiple cell types, for nearly all pathways examined: Wnt, Notch, Hedgehog, BMP, VEGF, FGF, semaphorins and ephrin in Rgl1-3; CNTF in ependymal cells; TNF in microglia, IGF and PDGF in pericytes, and TGFβ in endothelial cells. No evidence for Slit-Robo and endothelin autocrine signaling was found, but rather for possible directional paracrine signaling from Rgl3 to endothelial cells (Slit-Robo) or from endothelial cells and microglia to Rgl1-3, ependymal and pericytes (endothelin) (Fig. 4B, C). Combined, our findings identify radial glia cells, in particular Rgl3, as the midbrain-specific cell type that expresses the most complete combination of ECM components and ligands for signaling pathways in midbrain development.

### Transcriptional networks in Rgl2 and Rgl3

We previously identified floor plate radial glial cells as mDA progenitors (Bonilla et al., 2008). However, our recent identification of three different types of radial glial cells in the ventral midbrain (La Manno et al., 2016) raised the question of which of the three identified radial glia cell types is the mDA progenitor that undergoes mDA neurogenesis. We thus decided to examine the transcription factors expressed in each radial glia cell type using curated databases with identified target genes (Janky *et al*, 2014). Transcription factors in the database were filters to include only those expressed by Rgl1-3, and then we clustered them according to shared target genes (Fig. 5A, EV6A-B). We next investigated which of the possible transcription factor combinations can best explain the transcriptional profile of Rgl1-3 (Fig. 5B). Analysis of the statistical significance of the multiple combinations of transcription factors that up-regulate target genes in Rgl1-3 was performed using the Fast Westfall-Young (FWY) permutation test for multiple testing correction (Terada *et al*, 2013b). This procedure also prunes for non-significant combinations (Tarone, 1990), transforming the combinatorial analysis into a tractable problem. The specificity of the transcription factor combination analysis was confirmed by performing a control FWY analysis of transcription factors randomly selected from MSigDB while maintaining the transcriptional profile of each cell type. Random selections of transcription factors in Rgl1-3, resulted in much fewer significant combinations than obtained with the transcription factors expressed in each cell (Fig. EV6C). Thus, our results show that it is possible to obtain insights of the transcriptional networks responsible for the transcriptome of a cell type by integrating information about transcription factor binding sites and individual cell type transcriptomes.

**Figure 5.**
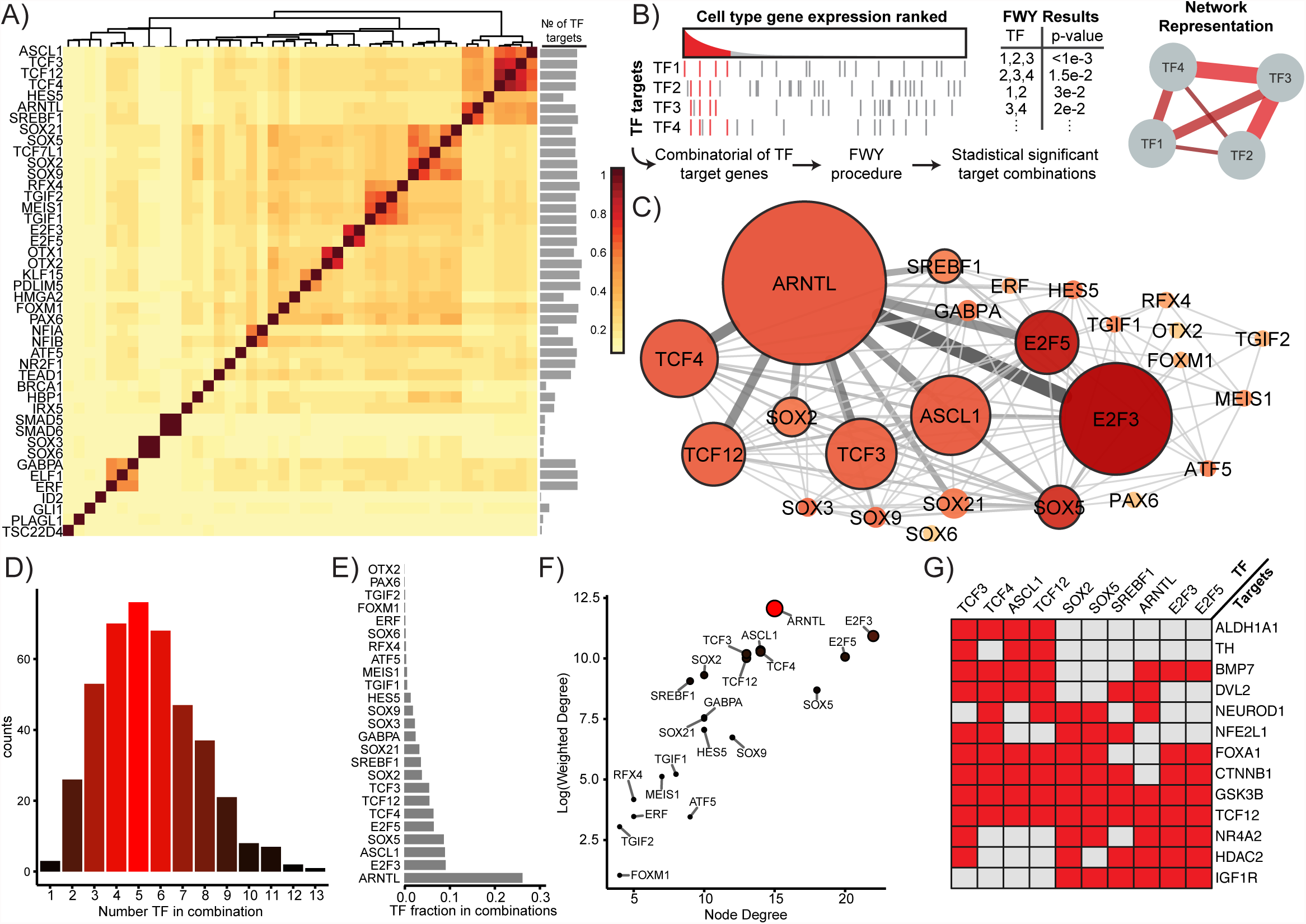
Combinatorial analysis of the enrichment of transcription factors in Rgl1. **(A)** Clustering of transcription factors expressed in Rgl1 by Jaccard index of shared target genes. A bar plot to the right represents the number of targets genes for each transcription factor. **(B)** Diagram of the combinatorial analysis of transcription factors (TF) by their enrichment in common targets genes. Left, enriched genes are those regulated by TF1-4 and expressed above threshold. Red lines represent upregulated target genes. Several target genes were shared by different TF. Right, network representation of TF sharing target genes, analyzed by Fast Westfall-Young (FWY) multiple combinatorial transcription factor enrichment. **(C)** Network representation of FWY analysis result of Rgl1. Node color is proportional to node degree and its size is proportional to weighted degree. Nodes with high weighted degrees (core nodes) have a black border. Color intensity and width of the line connecting two nodes (edge) is proportional to the interaction score of the TF pair. (**D)** Distribution of number of TF involved in significant combinations. **(E)** Frequency distribution of significant TF permutations distributed accordingly to their abundance. **(F)** Plot comparing node degree and weighted node degree of the network. For presentation purpose, weighted degree is shown in logarithmic scale. Color intensity and size is proportional to weighted degree. **(G)** Heat map of selected target genes of core transcription factors enriched on Rgl1 (red) found in databases.

Analysis of Rgl2 identified two large transcription factor clusters composed by Sox and proneural basic helix-loop-helix (bHLH) transcription factors (Fig. EV6A). We next investigated by which of the possible transcription factor combinations can best explain the transcriptional profile of Rgl2. However, FWY analysis showed that the only significant combination of transcription factors in Rgl2 was that formed by *PAX6* and *TCF7L1* (p-value=0.025), which has been reported to maintain neural progenitors in undifferentiated state (Kuwahara *et al*, 2014). Analysis of the target genes for these two transcription factors reveled enrichment of GO terms associated to development of the forebrain (p-value: 3.12e-45), hindbrain (p-value: 7.88e-8) and midbrain (p-value: 5.89e-5), suggesting a generic role in maintenance of neural progenitors. Moreover, Rgl2 cells and PAX6 expression are found in the basal plate of the midbrain, a compartment that does not give rise to mDA neurons but is rather involved in glial cell development (La Manno *et al*, 2016).

Examination of Rgl3 also identified two large clusters formed by Sox and bHLH genes (Fig. EV6B). FWY analysis of Rgl3 revealed 15 enriched transcription factor combinations, which formed a network centered on *TEAD1*, a component of the hippo pathway (Harvey & Hariharan, 2012), as well as *RFX4, PDLIM5* and *SOX13* (Fig. EV6D-E). Analysis of the targets genes of these transcription factors, identified the IGF and Wnt signaling pathways, followed by the regulation ECM components (Fig. EV6F), which again underlines a role of Rgl3 in the formation of the mDA niche.

### Identification of a transcriptional network in Rgl1 involved in mDA neurogenesis

We next analyzed Rgl1, the radial glia that appears the earliest in the developing VM (Fig. 3E) as well as the one expressing less ECM (Fig. 3A) and more receptors than ligands for developmental pathways (Fig. 4A). As it was the case for Rgl1 and Rgl3, two large Sox and bHLH transcription factor clusters with common target genes were identified (Fig. 5A). However, our FWY analysis revealed that a total of 25 transcription factors generate 419 significant combinations (Table EV4). We used a network to represent the combinations of transcription factors. A pairwise interaction score was calculated for all significant transcription factor combinations, based on the frequency and p-value (Fig. 5C). The mean number of transcription factors per combination was 5.465 (Fig. 5D) and the five most common transcription factors were *ARNTL*, *E2F3*, *ASCL1*, *SOX5* and *TCF4* (Fig. 5E). The transcription factors E2F5 and E2F3, two cell cycle regulators (Chen *et al*, 2009; Trimarchi & Lees, 2002), showed the highest number of interaction partners (node degree, Fig. 5F), but these interactions were less frequent and with higher p-values. When node degrees were weighted for interaction score, *ARNTL* was identified as the central gene in the network (Fig. 5F), followed by *E2F3, ASCL1*, *E2F5, TCF3*, *TCF4*, *TCF12*, *SOX2, SOX5* and *SREBF1*. These results indicate that components of the bHLH neurogenic cluster, such as *ARNTL, ASCL1, TCF3,TCF4 and TCF12,* explain most aspects of the transcriptional state of Rgl1. Moreover, some of the genes in this cluster have been previously found to regulate mDA neurogenesis, such as *Ascl1,* which works in concert with *Neurog2* (Kele *et al*, 2006); or TCF3 and 4, that interact with active β-catenin to control mDA neurogenesis (Arenas, 2014); or *SREBF1,* a direct target of the nuclear receptors *Nr1h2* and *Nr1h3* (Schultz *et al*, 2000), that are required and sufficient for mDA neurogenesis (Sacchetti *et al*, 2009; Theofilopoulos *et al*, 2013).

Target genes of the transcription factors in this network were analyzed for enrichment using the curated gene sets of the molecular signature database repository (MSigDB, (Subramanian *et al*, 2005)). We found that the main functions controlled by this network were the regulation of cell cycle as well as Wnt and Notch signaling (Fig. EV6G), all of which are fundamental for mDA neurogenesis (Louvi & Artavanis-Tsakonas, 2006; Arenas *et al*, 2015). Analysis of putative target genes of this network included Wnt signaling components such as *DVL2*, *CTNNB1*, *GSK3B* and *TCF12* (Fig. 5G) (Inestrosa & Arenas, 2010; Willert & Jones, 2006; Neuman *et al*, 1993). *FOXA1,* involved in Shh signaling and mDA neuron development (Hynes *et al*, 1995), as well as components of other signaling pathways such as *IGF1R* (Quesada *et al*, 2007) and *BMP7* (Brederlau *et al*, 2002). Genes involved in neurogenesis, such as *NEUROD1* (Cho & Tsai, 2004), *HDAC2* (Jawerka *et al*, 2010) and *TCF12* (Uittenbogaard & Chiaramello, 2002). As well as the transcriptions factors required for mDA neuron development, such as *NFE2L1* (Villaescusa *et al*, 2016) and the nuclear receptor *NR4A2* (Zetterström *et al*, 1997), as well as markers of mature mDA neurons, *TH* and *ALDH1A1* (Arenas *et al*, 2015), were also found. Thus our results indicate that a transcriptional network mainly formed by *ARNTL, TCFs* (*3*, *4* and *12*), *ASCL1*, *SOXs* (*2* and *5*) and *SREBF1* can set in motion a transcriptional program allowing them to control cell cycle exit and neurogenesis, respond to Wnt and Shh signaling and differentiate into postmitotic neuroblasts and mDA neurons.

Combined, our analysis of transcription factor networks and their target genes suggest that each Rgl cell type contribute to diverse aspects of VM and mDA neuron development. While Rgl2 expresses transcription factors known to control progenitor maintenance, Rgl3 expresses niche signals. Notably, the only radial glia cell type found to expresses a significant combination of transcription factors known to regulate mDA neurogenesis was Rgl1, suggesting that this is the cell type previously found to undergo mDA neurogenesis in the VM (Bonilla et al., 2008).

### ARNTL is required for mDA neurogenesis

Our analysis of Rgl1 identified ARNTL as the central node and the most predominant transcription factor in a network that explains the transcriptional profile of Rgl1. *ARNTL* is known to promote neurogenesis, control cell cycle exit and regulate circadian rhythm (Bouchard-Cannon *et al*, 2013; Kimiwada *et al*, 2009; Malik *et al*, 2015). However, it is at present unknown whether it plays any role in VM development or mDA neurogenesis. We thus decided to examine whether *ARNTL* is both sufficient and required for mDA neuron development. For this purpose, we first examined the presence of ARNTL in the developing midbrain and found that it is present in Sox2^+^ cells in the developing midbrain floor plate at E13.5 (Fig S7A). To test its functionality we then took advantage of a human long-term neuroepithelial stem (hLT-NES) cell line derived from human embryos (Sai2, Tailor *et al*, 2013), and performed *ARNTL* gain and loss of function experiments (Fig. 6A,B, S7B) during mDA neuron differentiation (Villaescusa *et al*, 2016). As previously reported, hLT-NES cells acquired a midbrain floor plate phenotype, assessed by the presence of both FOXA2 and LMX1A (Fig. 6C) and immunoreactivity for DA markers such as TH and NR4A2 at day 8 of differentiation (Fig. 6D).

**Figure 6.**
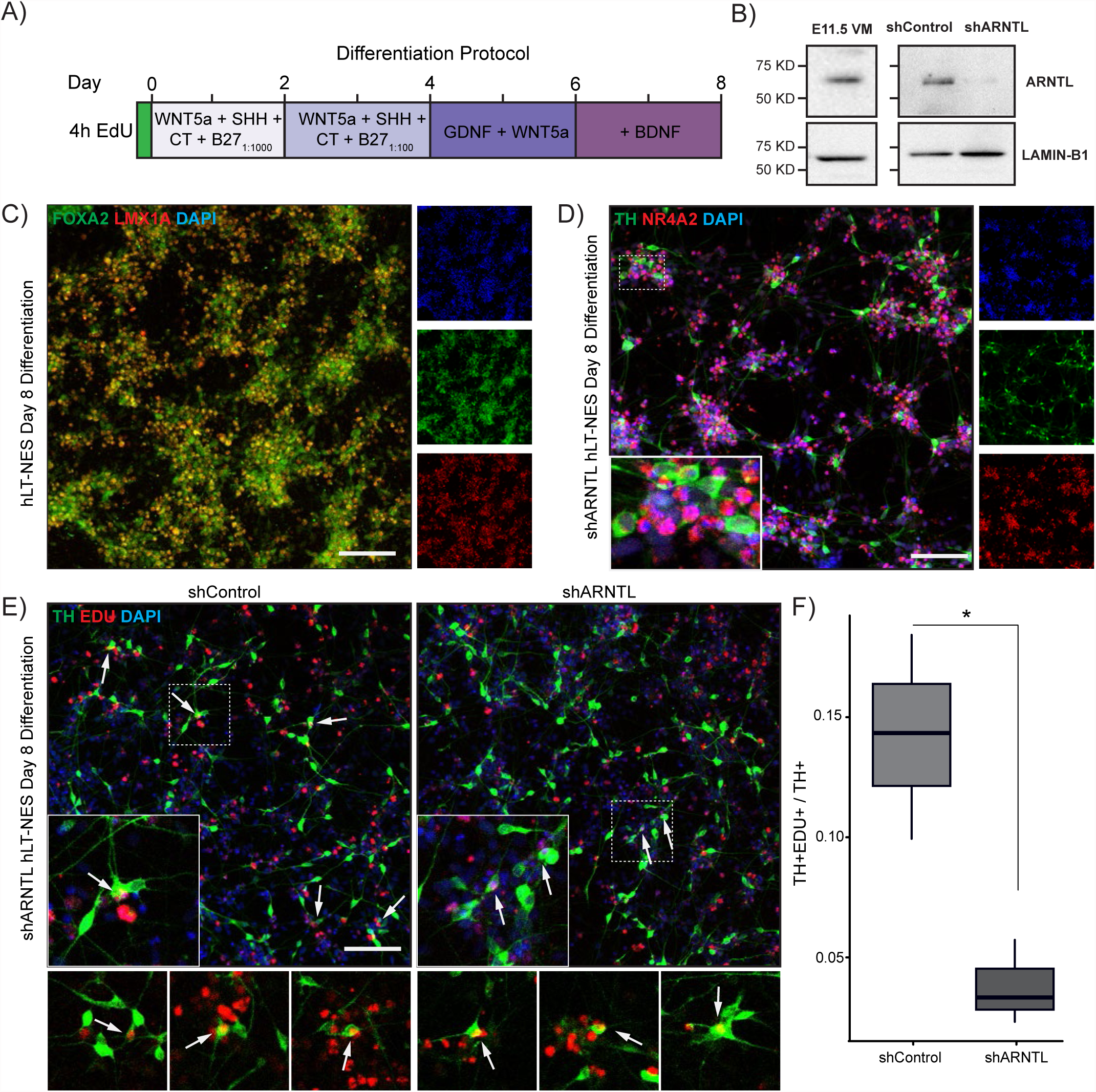
ARNTL is required for the differentiation of human neuroepithelial stem cell into mDA neurons. **(A)** Representation of the protocol to examine dopaminergic neurogenesis in hLTNES cells. **(B)** Western blot analysis identified the presence of ARNTL in the mouse ventral midbrain at E11.5 (left) and in control hLT-NES cells. ARNTL was dramatically reduced by shRNAs against ARNTL (right). LAMIN-B1 was used as loading control. **(C)** The midbrain floor plate marker FOXA2 (green) and the DA lineage marker LMX1A (red) are present in hLT-NES at day 8 of differentiation. (D) hLT-NES differentiated for 8 days are immunoreactive for the DA markers TH (green) and NR4A2 (red). **(E)** Analysis of neurogenesis in shControl and shARNT hLT-NES cells as identified by the presence of cells double positive (arrows) for TH (green cytoplasm) and EdU (red nuclei). **(F)** Quantification of mDA neurogenesis: TH^+^ and EdU^+^ cells relative to the total TH+ cells. (p-value = 0.0362, N = 3). Scale bars in C, D and E, 100 μm.

For gain of function of experiments, proliferating hLT-NES cells were transduced at day -2 with a lentivirus carrying a tetracycline-inducible vector (Gossen *et al*, 1995) encoding for ARNTL, or GFP as control. Cells were treated with doxycycline from day-1 to day 1 and given a 4 h pulse of EdU before day 0, to mark proliferating cells that undergo mDA neurogenesis and become EdU^+^ and TH^+^ at day 8 after differentiation (Fig S7B). Analysis of ARNTL levels at day 0 revealed a 75% increase in protein levels compared to endogenous levels in GFP-expressing control cells (Fig. S7C). Notably, we found that *ARNTL* overexpression at this level did not affect the proportion of TH^+^ cells undergoing neurogenesis (double EdU^+^ and TH^+^, Fig. S7D, E), but higher levels of ARNTL impaired their survival. These results suggested that ARNT overexpression “*per se*” is either not active in this process or that it cannot increase mDA neurogenesis beyond what endogenous levels ARNTL already achieve in Lt-NES cells. We thus examined whether endogenous ARNTL is required for mDA neurogenesis.

We next performed loss of function experiments using lentiviral vectors to stably knockdown endogenous ARNTL in hLT-NES cells. The resulting cell line, shARNTL-hLT-NES, exhibited a dramatic reduction of ARNTL protein levels, compared to a control shRNA (shControl, Fig. 6B). As with the gain of function, hLT-NES cells were pulsed with EdU before differentiation, to examine neurogenesis. Analysis of cells differentiated for 8 days revealed that the percentage of EdU^+^ and TH^+^ cells was reduced by 80%, from 14.23 ± 0.04% (mean ± SD) in shControl to 3.79 ± 0.017% in shARNTL-hLT-NES (Fig. 6E, F). Thus, our results show that endogenous ARNTL is *required* for DA neurogenesis and support the emerging concept that a network of bHLH transcription factors controls this process.

## DISCUSSION

In this study, we have gained a broad view of the transcriptional programs operating in the VM and at single-cell level during mDA neurogenesis. Our analysis identifies a transcriptional program that defines mDA neuron development and Rgl1-3 as the main cell types expressing genes that regulate mDA neurogenesis and the mDA niche, providing thus novel insights into midbrain development at a single-cell level. Notably, we found that Rgl1 selectively expresses transcription factors controlling target genes required for mDA neurogenesis and cell membrane receptors for multiple developmental signals. On the other hand, Rgl2 was found to express transcription factors responsible progenitor maintenance, receptors for developmental factors and genes involved in lipid metabolism. Finally, Rgl3 was found to express transcription factors regulating the formation and maintenance of the mDA niche, as well as multiple ECM components and ligands controlling several developmental pathways (Fig. 6). Thus, our study suggests that each of the three radial glia cell types (Rgl1-3) contributes to different aspects of mDA neuron development.

Our analysis of the VM transcriptome identified a mDA module that was mainly contributed to by seven cell types: Rgl1-3, ependymal cells, endothelial cells, pericytes and microglia. However, the most abundant cells contributing to mDA neurogenesis were Rgl1 (E11.5-E12.5), Rgl2 (E13.5-E14.5) and Rgl3 (E15.5). Moreover, two of them, Rgl1 and Rgl3, are present in the midbrain floor pate (La Manno *et al*, 2016), the anatomical compartment that generates mDA neurons (Arenas *et al*, 2015). While both cells contribute to the expression of developmental signals that control mDA neurogenesis, we found that Rgl3 is the main contributor to ligands and ECM. We also identify Rgl1 as the main cell type expressing receptors for midbrain developmental signals. Thus, our data suggest a model in which Rgl1 controls neurogenesis in an autocrine manner from E11.5 to E13.5 and Rgl3 in a paracrine fashion from E13.5-E14.5. This switch in neurogenic regulation may be of importance for mDA neuron development as *substantia nigra* mDA neurons emerge earlier than those in the ventral tegmental area. We thus examined the expression of ligands in early radial glia (Rgl1) versus late radial glia (Rgl3) for pathways whose receptors are expressed in Rgl1. While no change in expression was detected for ligands such as *Shh, Sema3b, Sema5b, Efna4* and *Efnb1*, a switch was detected from *Wnt7a and Wnt7b* in Rgl1 to *Wnt5a and Wnt5b* in Rgl3. In addition, other factors were expressed in early Rgl1, such as *Jag1*, or in late Rgl3, such as *Sfrp1, Dkk3, Bmp1, Fgf7, Ntn1, Dcn* and *Spon1*. These results suggest that one of the major changes taking place in early vs late mDA neuron niche is a switch form early Wnt/β-catenin signaling by Wnt7a (Fernando *et al*, 2014) to late Wnt/β-catenin-independent signaling by Wnt5a (Andersson *et al*, 2008). This switch was further confirmed by the late action of two pro-differentiation Wnt/β-catenin inhibitors such as Sfrp1 (Kele *et al*, 2012) and Dkk3 (Fukusumi *et al*, 2015), reinforcing thus a role for Wnt/β-catenin-independent signaling at late stages. The necessity of an appropriate balance between these two pathways has been recently confirmed by analysis of double *Wnt1* and *Wnt5a* mutant mice (Andersson *et al*, 2013). Our study thus extends to Wnt5b and Wnt7b the spectra of candidate Wnt ligands participating in this complex balance and identifies the precise cell types expressing Wnts and their receptors during mDA neuron development.

We also found that cell types that are not midbrain specific, such as endothelial cells, pericytes and microglia, express factors relevant for mDA neurogenesis, such as *Wnt5b* and *Jag1* (Fig. 4A). Notably, the vasculature is known to be in contact with radial glia cells and their process during development, allowing thus for direct or indirect modulation of mDA neurogenesis. We thus suggest that these cells can be part of a generic neurogenic niche, that may add on to midbrain-specific signals derived from radial glia.

Our bioinformatics analysis of the transcription factor networks in Rgl1-3 also suggest individual functions for each radial glia cell type. These range from controlling neurogenesis (Rgl1), to progenitor maintenance (Rgl2) and niche formation (Rgl3). Notably, our results suggest that the radial glia previously identified to undergo mDA neurogenesis (Bonilla *et al*, 2008), is Rgl1. Indeed, Rgl1 expressed a neurogenic bHLH network formed by *Arntl, Ascl1*, *Tcf3*, *4*, *12* and *Srebf1*. Previous studies have linked transcription factors such as *Ascl1* to mDA neurogenesis, via *Neurog2* (Kele *et al*, 2006), or *Tcf3/4* regulated by Wnt/β-catenin signaling (Arenas, 2014) or *Srebf1* (Schultz *et al*, 2000) by *Nrh2*-*3/Lxr* (Sacchetti *et al*, 2009; Theofilopoulos *et al*, 2013). Moreover, the Rgl1 network also includes two members of the Sox family of transcription factors. *Sox2*, which is expressed in floor plate progenitors (Kele *et al*, 2006; Bonilla *et al*, 2008) and *Sox5*, which promotes cell cycle exit and differentiation (Martinez-Morales *et al*, 2010). Notably, this network was centered on the clock gene *ARNTL*, also known as *Bmal1*, a pioneer transcription factor (Menet *et al*, 2014). ARNTL controls cell cycle entry/exit and neurogenesis (Bouchard-Cannon *et al*, 2013; Malik *et al*, 2015) and directly regulates the expression of *Neurod1* (Kimiwada *et al*, 2009), a proneural transcription factor that initiates neuronal differentiation and migration (Peyton *et al*, 1996; Pataskar *et al*, 2015). *ARNTL* had not been previously linked to mDA neurogenesis, but based on key role predicted by our network analysis, we investigated the role of this transcription factor and found it is required to promote mDA neurogenesis. Thus, our results identify ARNTL as a novel factor controlling mDA neuron development, and support the idea that several bHLH transcription factors control mDA neurogenesis.

In sum, our study examined the transcriptome of the developing VM during mDA neurogenesis and identifies two radial glia cell types, Rgl1 and Rgl3, as the main components of the mDA neurogenic niche. While Rgl1 expresses transcription factors and target genes required for mDA neurogenesis, Rgl3 expresses core ECM components and most ligands required to control midbrain development and mDA neurogenesis. These cells are part of an extended niche formed by a neural component (Rgl1-3 and ependymal cells), a non-neural component (endothelial cells and pericytes and microglia). Our results thus uncover the diversity and richness of the molecular components of the mDA niche, their cellular origin and their temporal dynamics during mDA neurogenesis. Moreover, our results identify novel components of the mDA niche, including a novel gene required for mDA neurogenesis, *ARNTL*, and provide new knowledge that can be applied to improve current regenerative medicine approaches for the treatment of Parkinson’s disease.

## AUTHOR CONTRIBUTIONS

EMT performed the biological validation experiments. EMT and GLM performed the data analysis. GLM and SI performed the RNA extraction and cDNA library preparation. GLM ran the NGS pipeline. DG and CV performed the embryonic dissections. PRdVC performed the preparation of lentiviral particles. SL helped to design the experiments and interpreted results. EMT and EA designed experiments, interpreted results and wrote the manuscript. All authors have read and approved the manuscript.

## ACKNOWLEDGMENTS

We thank members of the Arenas lab for their help and suggestions; and Johnny Söderlund and Alessandra Nanni for technical and secretarial assistance. Knut and Alice Wallenberg Foundation for support to the CLICK imaging facility at KI. Financial support was obtained from Swedish Research Council (VR projects: DBRM, 2008:2811, 2011-3116 and 2011-3318), Swedish Foundation for Strategic Research (SRL program), European Commission (NeuroStemcellRepair and DDPD-Genes), Karolinska Institutet (SFO Thematic Center in Stem cells and Regenerative Medicine) and Hjärnfonden to EA, grants from the Swedish Research Council and EU - FP7 program DDPDGENES to S.L (www.ddpdgenes.eu) and a fellowship from the Swedish Research Council to E.M.T.

## MATERIALS AND METHODS

### Bulk transcriptome determination

Mouse embryos were obtained from TH-GFP animals (Matsushita *et al*, 2002) that were mated overnight, and noon of the day the plug was considered E0.5. Embryos were dissected out of the uterine horns at E11.5-E14.5 and placed in ice-cold sterile PBS where brain regions were dissected under a stereomicroscope with a UV attachment to detect GFP. VM samples corresponded to domains M3 to M7 of the floor and basal plate (Nakatani *et al*, 2007). Tissue samples were collected in separate tubes, stored at -80°C until RNA isolation. Ethical approval for mice experimentation was granted by the local ethics committee, Stockholm Norra Djurförsöksetiska Nämnd number N326/12 and N158/15.

Total RNA isolation was performed with RNeasy kit (Qiagen). The RNA integrity and concentration was checked using Qubit and 2200 TapeStation (Agilent). Illumina TruSeq libraries were prepared using kit and protocols from Illumina. High-throughput sequencing was performed on a HiSeq 2000.

### Genes Expression Analysis

Differentially expressed genes (DEG) on each developmental stage were identified utilizing Qlucore Omics Explorer v3.1 (Qlucore AB, Lund, Sweden) utilizing t-test comparing VM samples versus other regions, with false discovery rate q-value correction (Storey, 2002). Variance filter of 15% and a threshold for significance of p-value = 0.01 across stages, unless otherwise stated. Sample correlation was calculated from log2(RPKM+1) values after variance filtering of 12.5%. Single-cell expression data was obtained from La Manno et al. 2016. Significant expression is baseline (>99.8% posterior probability). Analyses were made using R (https://www.rproject.org) and ggplot2 for plots (http://ggplot2.org/).

### Weighted Gene Co-Expression Network Analysis (WGCNA)

(Langfelder & Horvath, 2008) was performed with the log2(RPKM+1) transformed values, filtered by variance until 10,068 genes were selected. The topological overlap matrix was calculated with the variables of a signed network, with power of 7. The identification of modules was performed with the “tree” option on a minimum module size of 100; modules with correlation higher than 90% were merged. Module enrichment for DEG in VM at all analyzed stages was done with Fisher’s exact test with false discovery rate q-value correction (Storey, 2002). A score for the enrichment was calculated as the product between the −log10(q-value) and a standardized z-score for DEG per module. Module network layout was made using Cytoscape and interactions in the top 5% of adjacency were selected for further analysis. Expression changes over time are represented by the color of each node, which was calculated as the normalized difference of RPKM between late (E13.5, E14.5) and early (E11.5, E12.5) for each gene.

The identification of VM gene modules with developmental or stage dependent expression were identified with WGCNA on the VM samples from E11.5 to E14.5. Further filtering for genes not expressed in the VM was performed until 9,061 genes were selected. The topological overlap matrix was calculated with the options of a signed network, with power of 17. The identification of modules was performed with the “tree” option on a minimum module size of 100 genes, modules with correlation higher than 99% were merged. Correlation with samples trait and Student asymptotic p-values were calculated as described (Langfelder & Horvath, 2008). Embryonic stage (E11.5 to E14.5) was used as sample trait, ordinal values for stage (1 for E11.5, 2 for E12.5 and so on), and binary values for middle stages (E12.5, E13.5 as 1). Network layouts and analysis, were made using Cytoscape v3.3.0 (Shannon *et al*, 2003) or Gephi v0.9.1 (Bastian *et al*, 2009). WGCNA were carried out with the R package v1.49 (Langfelder & Horvath, 2008).

### Score for cell types contribution to ECM or signaling

Cell type contribution for ECM or signaling were calculated with the data from single cell of mouse VM (La Manno *et al*, 2016). The identified transcripts corresponded to cells found in the floor plate and basal plate of the midbrain, domains M3 to M7 (Nakatani *et al*, 2007). We selected the two components for each analysis. ECM was the result of the ECM core genes and ECM regulation gene sets (Naba *et al*, 2012). For signaling, curated ligands and receptors gene sets (Table EV3).

The score for a gene set is calculated as:

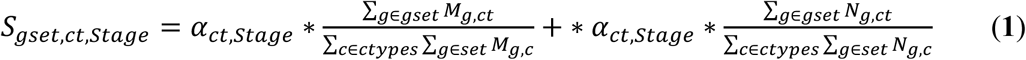

Let *M_g,ct_* be the Bayesian estimate of the expression level of the gene *g* in cell type *ct* and let *N_g,ct_* be the value of the indicator vector that is 1 when the gene *g* is expressed in cell type *ct* above baseline and 0 otherwise. Let *α_ct_* be the relative sampling ratio of cell type *ct* in a developmental stage, with a value of 1 when the analysis include all the developmental stages. The score is an indicator between diversity of genes in a biological function or gene set and the expression levels of those genes. A combined score by the sum of both components give us an ECM or signaling scores. This simplification of a biological process in a tissue, provide an indicator for a biological processes break down by the cell types found in that tissue. The threshold line was set at the percentile 99.9% of the bootstrapped distribution of the mean of the combined score with 10e5 replicates.

### Pathway receptor ligand

A list of ligand (159 genes) and receptors or co-receptor (114 genes) was curated from the literature, representing 21 signaling pathways. Ligands and receptors were grouped in pathways, without assuming specific ligand-receptor pair (Table EV3).

### Network single-cell deconvolution

Using data from La Manno et al., each gene in the network was assigned to the cell types that expressed that gene, as identified above baseline (see above and (La Manno *et al*, 2016)). Cell types represented by one gene, or cell types with a final contribution less than 1% of the genes were excluded from the network.

### Single-Cell Transcription Factor pattern mining

Transcription factors expressed on each radial glia cells were used as input to obtain the target of the human homologous from iRegulon plugin for Cytoscape (Janky *et al*, 2014), retrieving up to 1,000 target genes per transcription factor. Targets of transcription factors by cell type detailed on table EV5. Clustering of the transcription factor was done by Jaccard index, as an indicator of shared target genes.

Analysis of enrichment of combinatorial transcription factor target genes were done using Fast Westfall-Young (FWY) permutation procedure v1.0.1 (Terada *et al*, 2013a, 2013b). This is a computational efficient procedure for multiple testing correction with higher detection sensitivity than commonly used algorithms (Terada *et al*, 2013b). Upregulated or enriched genes among Rgl cells types was scored using the logarithmic difference between the groups (Subramanian *et al*, 2005). Values above 0.5 (or 50% of upregulation) were considered enriched in that cell type. Targets of the transcription factors were tested for enrichment in the upregulated genes in the one-tailed Fisher test with a significance threshold of 0.05, and tested with 1,000 permutations in the FWY to generate a null distribution of randomly permutated datasets (Terada *et al*, 2013b).

The resulting significant combinatorial patterns were represented as a network on which the edge between a set of transcription factors is quantified by an interaction score. This interaction score is calculated for each transcription factor pair as the sum of −log10 (adjusted p-value) for all the transcription factor combinations of that pair. Adjusted p-values <0.001 were consider equal to 0.001 for calculations of interaction score. Controls were performed with randomly selected transcription factors from MSigDB C3 database (Subramanian *et al*, 2005). Selecting the same numbers of transcription factors used before for Rgl1-3 respectively. For each cell type, 100 random transcription factors combinations were analyzed by FWY maintaining all other setting.

### Gene enrichment

Single-cell Gene Set Enrichment Analysis (GSEA) was done ranking the gene by the class difference for the cell type of interest. The analysis was performed on the GSEA software v2.2.2 (Subramanian *et al*, 2005) with the MSigDB genesets v5.0 for canonical pathways and GO biological processes (Subramanian *et al*, 2005). Due to the nature of the single-cell transcriptome profiles and molecular counting with unique molecular identifiers (Islam *et al*, 2014), negative enrichment score (ES) on a gene set for a cell type in particular have no meaning. Instead, negative ES were interpreted as enrichment on another compared cell type. Analysis of Gene Ontology (GO) enrichment were made utilizing the R package ClusterProfiler (Yu *et al*, 2012) or MSigDB (Subramanian *et al*, 2005) with false discovery rate q-value correction (Storey, 2002). Enrichment of the targets of transcription factor (Fig. EV6CF) were analyzed with hypergeometric test and with false discovery rate q-value correction (Storey, 2002) over the MSigDB C2 gene sets (Subramanian *et al*, 2005).

### Human neuroepithelial stem cell differentiation

Sai2 hLT-NES cells were maintained as described of hLT-NES cell (Tailor *et al*, 2013), DA differentiation and lentivirus infection were performed as previously described (Villaescusa *et al*, 2016). Each experimental condition was examined in three separate experiments and duplicate determinations.

Commercially available ARNTL shRNA lentiviral particles targeted against the human transcript were used (sc-38165, Santa Cruz Biotech.) to generate stable cell lines selected with puromycin (500 ng/ml first 4 days, maintained with 200 ng/ml). As negative control, lentiviral particles against no know human mRNA were used (sc-108080, Santa Cruz Biotech.). Expression levels of ARNTL were analyzed by western blot of total cell lysates, with anti-Arntl (1:1000, ab93806, Abcam). For loading control, anti-Lamin-B1 was used (1:5000, ab16048, Abcam). Differentiation experiments were done with stable shRNA cells with less than 10 passages counted from the lentiviral infection.

Mouse Arntl cDNA-pcDNA3.1 expression vector (Etchegaray *et al*, 2003) (kindly donate by Professors Steven Reppert and David Weaver, University of Massachusetts Medical School), was cloned into the lentiviral backbone Tet-O-FUW-EGFP (Addgene #30130) by blunt-end ligation into EcoRI blunted site of the backbone, replacing the EGFP open reading frame. The final plasmid was verified by sequencing before usage. Lentiviral production, infection and immunofluorescence were performed as previously described (Villaescusa *et al*, 2016; di Val Cervo *et al*, 2017). Inducible expression of ARNTL was done with doxycycline (Sigma, 50 μg/ml for day -1 and day 0, and 25 μg/ml for day 1).

Immunofluorescence was captured with a confocal microscope Zeiss LSM700 with a 10X 0.45NA objective, at a resolution of 0.3126 μm/pixel. A minimum of nine images per well were captured, with two technical replicates per condition in independent biological experiments (N=3 for LOF experiments, and N=4 for GOF experiments). Quantifications of EdU and DAPI positive nuclei were done utilizing Cell Profiler v2.2.0 (Jones *et al*, 2008). Double positive cell for TH and EdU, and total TH cells were counted manually in a condition-blind manner. This was done by randomizing each file name with *blindanalysis* perl script (Salter, 2016). Full images had linear levels adjusted for better visualization, done in Fiji (Schindelin *et al*, 2012).

## EXTENDED VIEW FIGURE LEGENDS

**Figure EV1.**
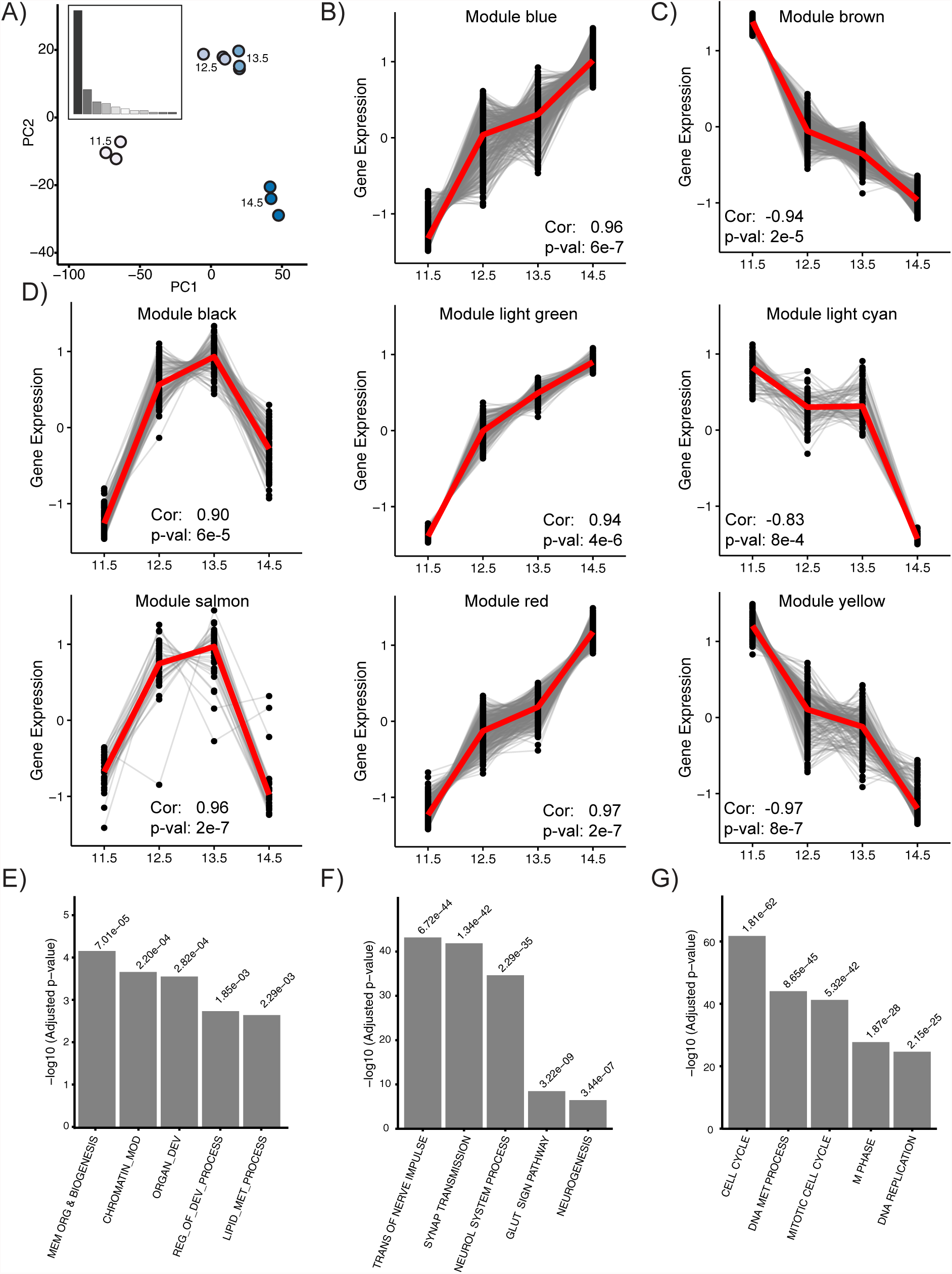
Ventral midbrain developmental genes modules. **(A)** Principal component analysis of VM samples at different developmental times. Insert shows the percentage of variance by component. **(B)** Modules with positive correlation of gene expression with development. **(C)** Modules with negative correlation to development. **(D)** Modules that correlate at E12.5 and E13.5. Correlations and p-values are detailed in each plot. **(E-G)** GO analysis of genes modules correlating at stages E12.5 and E13.5 **(E),** or with positive **(F)** or negative **(G)** correlation during development.

**Figure EV2.**
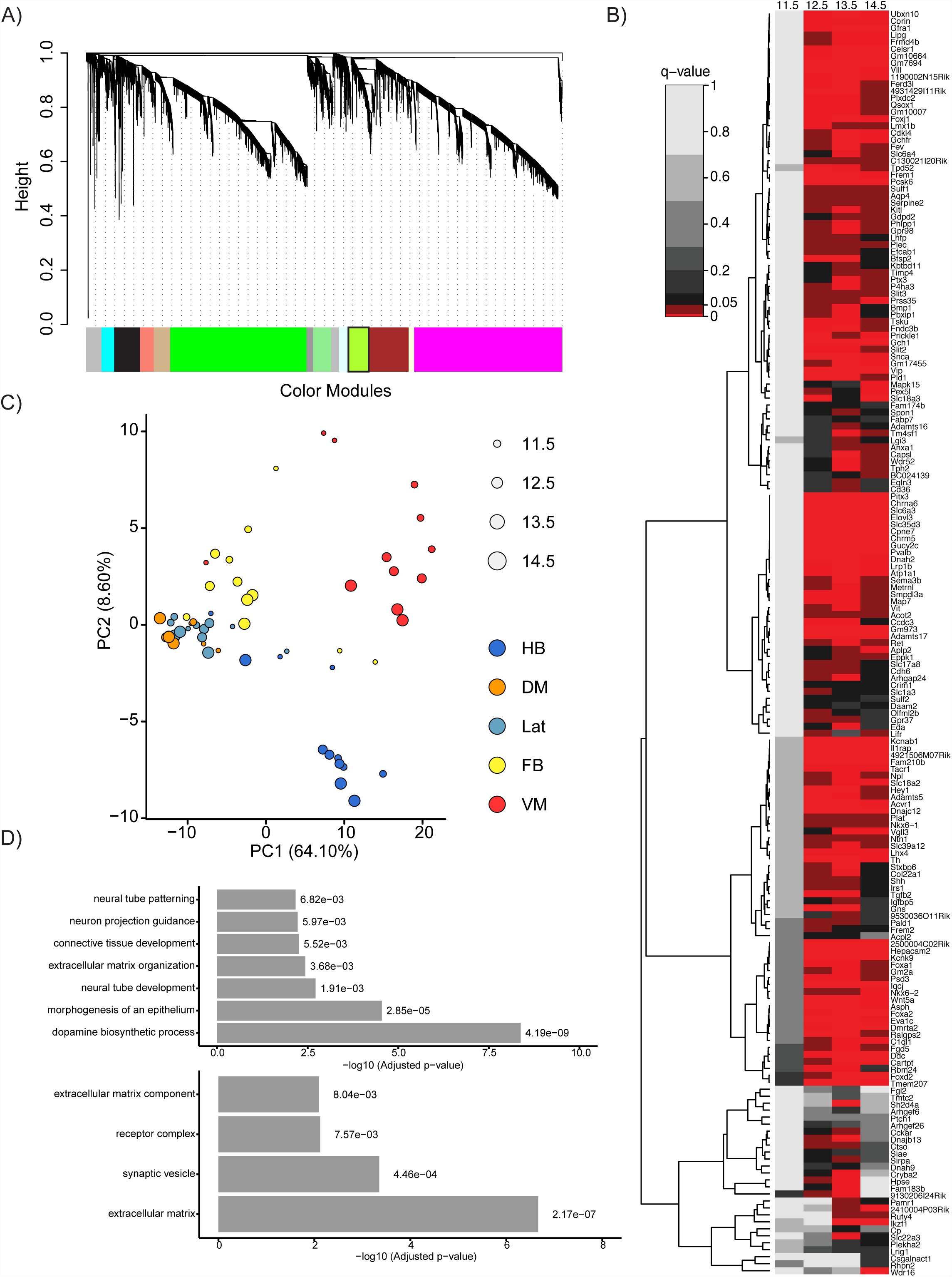
Weighted Gene Co-Expression Network Analysis of all samples. **(A)** Dendrogram of modules obtained from weighted gene co-expression network analysis. The mDA module is shown in “light green” and has black border. **(B)** Heat map of q-value of DEG in the mDA module for each developmental stage. Red scale, q-values between 0 and 0.05; black to light gray scale, q-values above 0.05. **(C)** Principal component analysis of the genes corresponding to the mDA module “light green”. The genes in the module were sufficient to cluster VM samples from the rest of the tissue samples. Color represents tissue and size represents developmental stage**. (D)** GO analysis of genes in the mDA module: Top, biological processes; Bottom, cellular components.

**Figure EV3.**
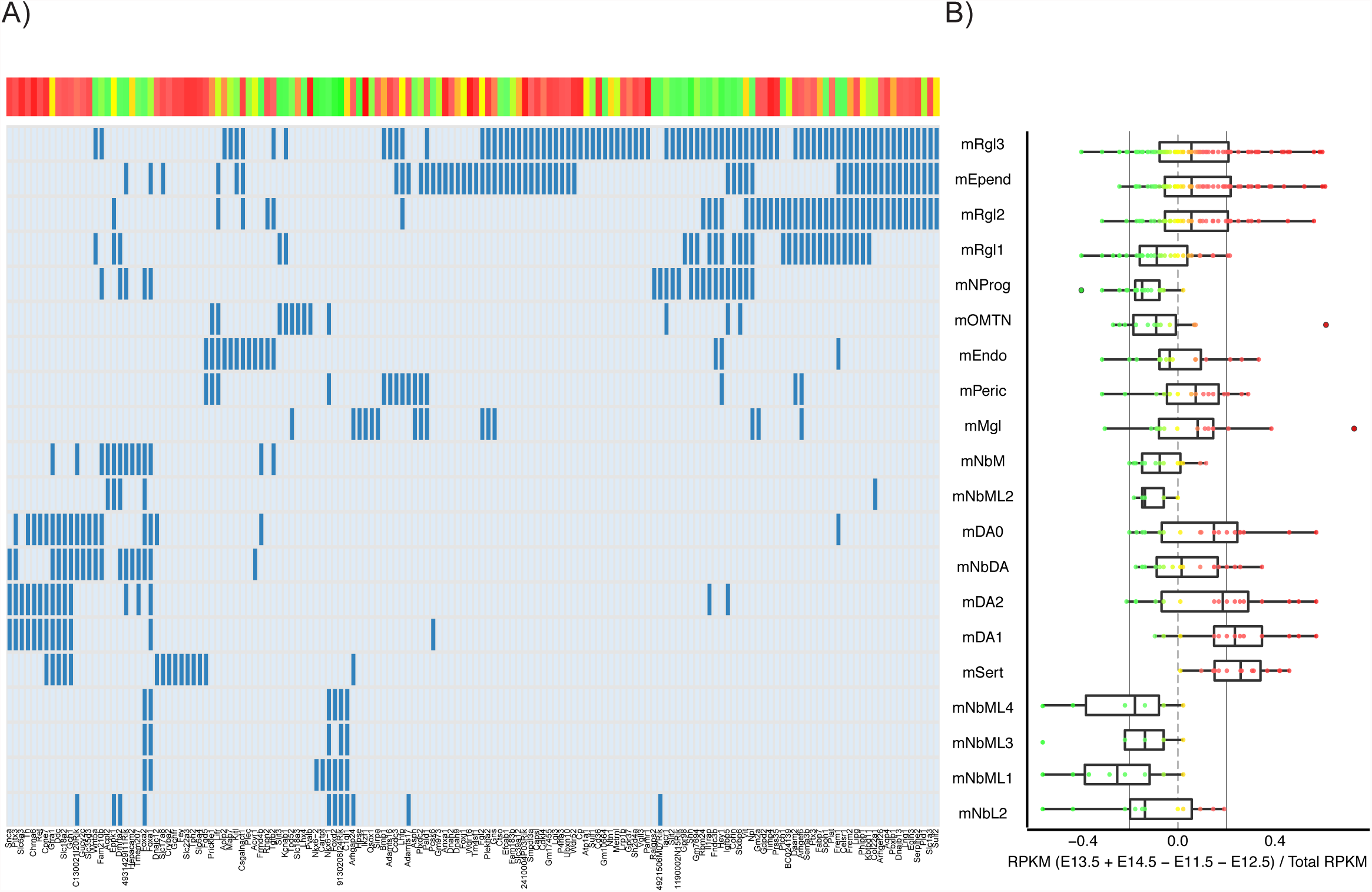
Single-cell network deconvolution. **(A)** Identity matrix of VM cell types and genes expressed in the mDA module. Blue lines represent significantly expressed genes in the corresponding cell type, compared to baseline. Red, gene increased; Yellow, maintained; Green, decreased over time. Color scale in figure 2a**. (B)** Analysis of gene expression levels (RPKM) in the mDA module for each cell type, comparing early (E11.5+E12.5) versus late stages (E13.5+E14.5).

**Figure EV4.**
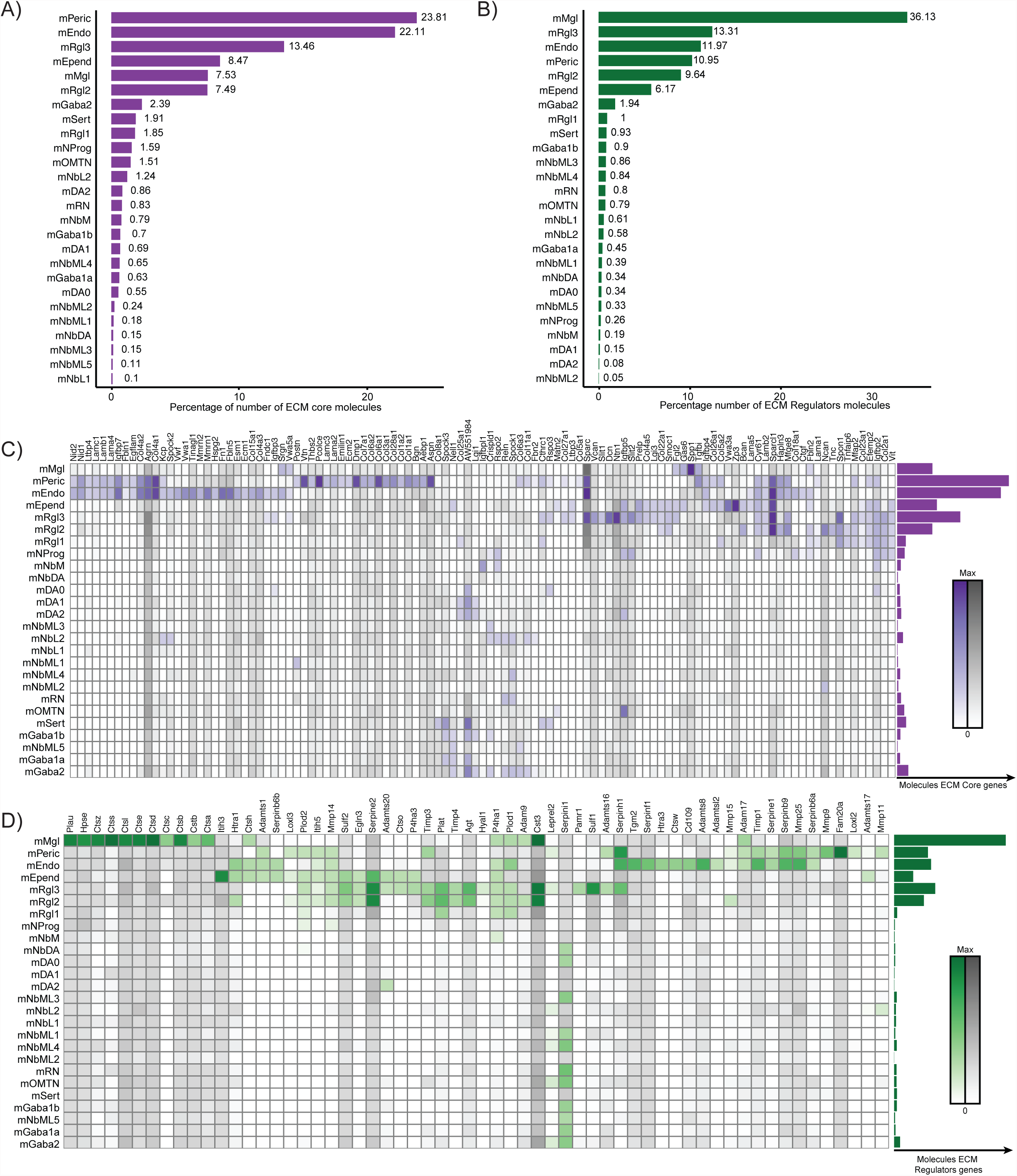
Contribution of different ventral midbrain cell types to the expression of extracellular matrix components and modifiers. **(A-B)** Bar plot with the contribution of each cell type to the total number of transcripts for core ECM genes **(A)** and regulatory ECM genes **(B). (C)** Heat map of ECM core component genes by cell type. Color intensity is proportional to the Bayesian estimate of expression level. Gray scale indicates values below significance level. Bar plot to the right represents total average of transcripts for ECM core components per cell type. **(D)** Heat map of the genes regulating ECM composition per cell type, represented as described above. Abbreviations: mMgl, microglia. mRgl1-3, radial glia type 1 to 3. mEndo, endothelial. mPeri, pericyte. mEpend, ependymal. mGaba1a, 1b and 2, GABAergic neurons 1a, 1b and 2. mSert, serotonergic neurons. mNbL1-2, lateral neuroblasts. mNbML1-5 mediolateral neuroblasts. mRN, Red Nucleus. mOMTN oculomotor and trochlear neurons. mNProg, neuronal progenitor. mNbM, medial neuroblast. mNbDA DA neuroblast. mDA0-2, DA neurons 1-2.

**Figure EV5.**
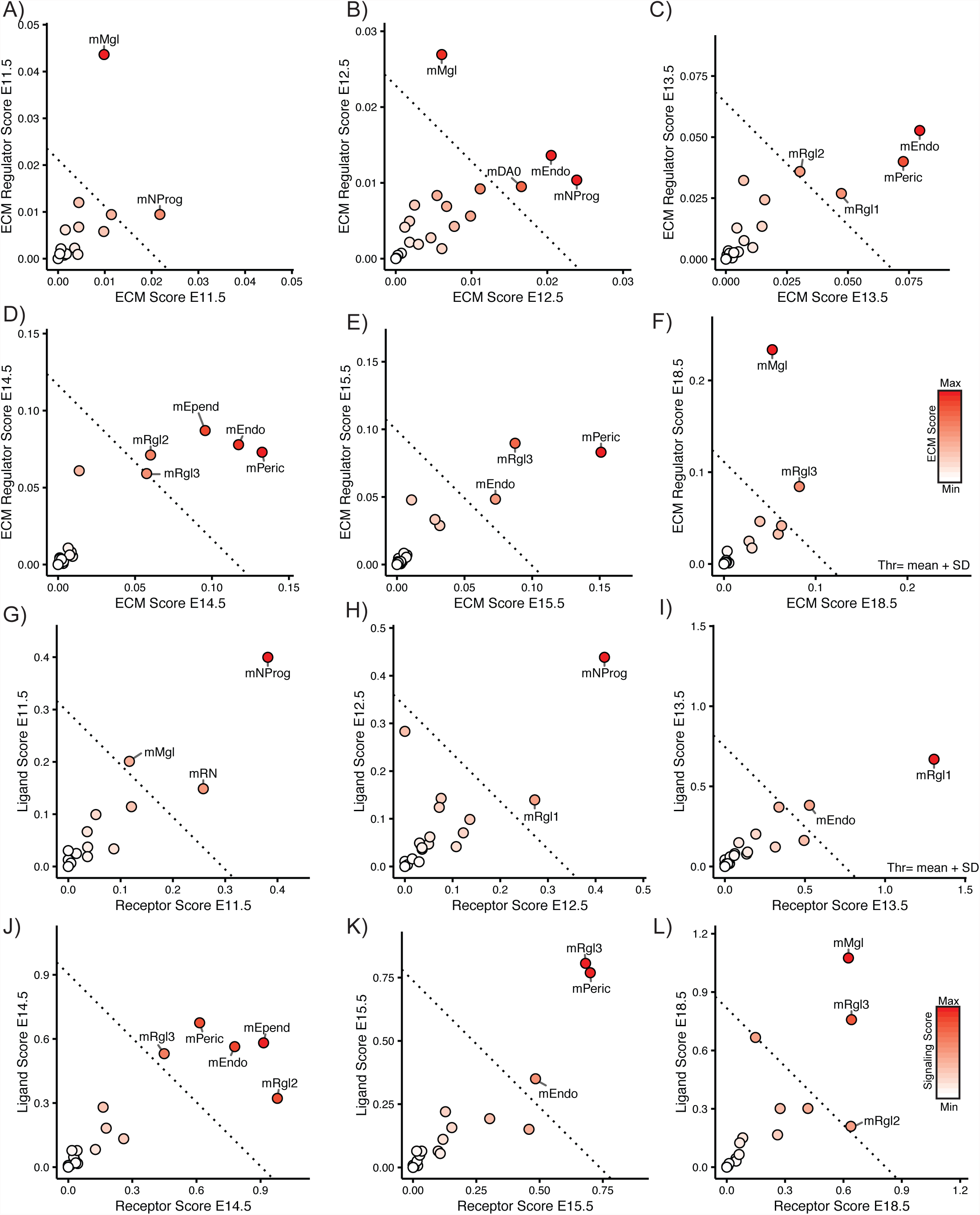
Stage dependent contribution of individual mouse VM cell types to signaling and ECM score in the mDA niche. **(A-F)** Plots showing the receptor and ligand scores of the different VM cell types considering cell type abundance at each developmental stages studied. **(G-L)** Plots showing the ECM regulator and ECM core score considering cell type abundance for each developmental stage. **(A, G)** E11.5, **(B, H)** E12.5, **(C, I)** E13.5, **(D, J)** E14.5, **(E, K)** E15.5 and **(F, L)** E18.5. Color intensity is proportional to the signaling score. Dotted line at mean plus standard deviation of the mean of the scores.

**Figure EV6.**
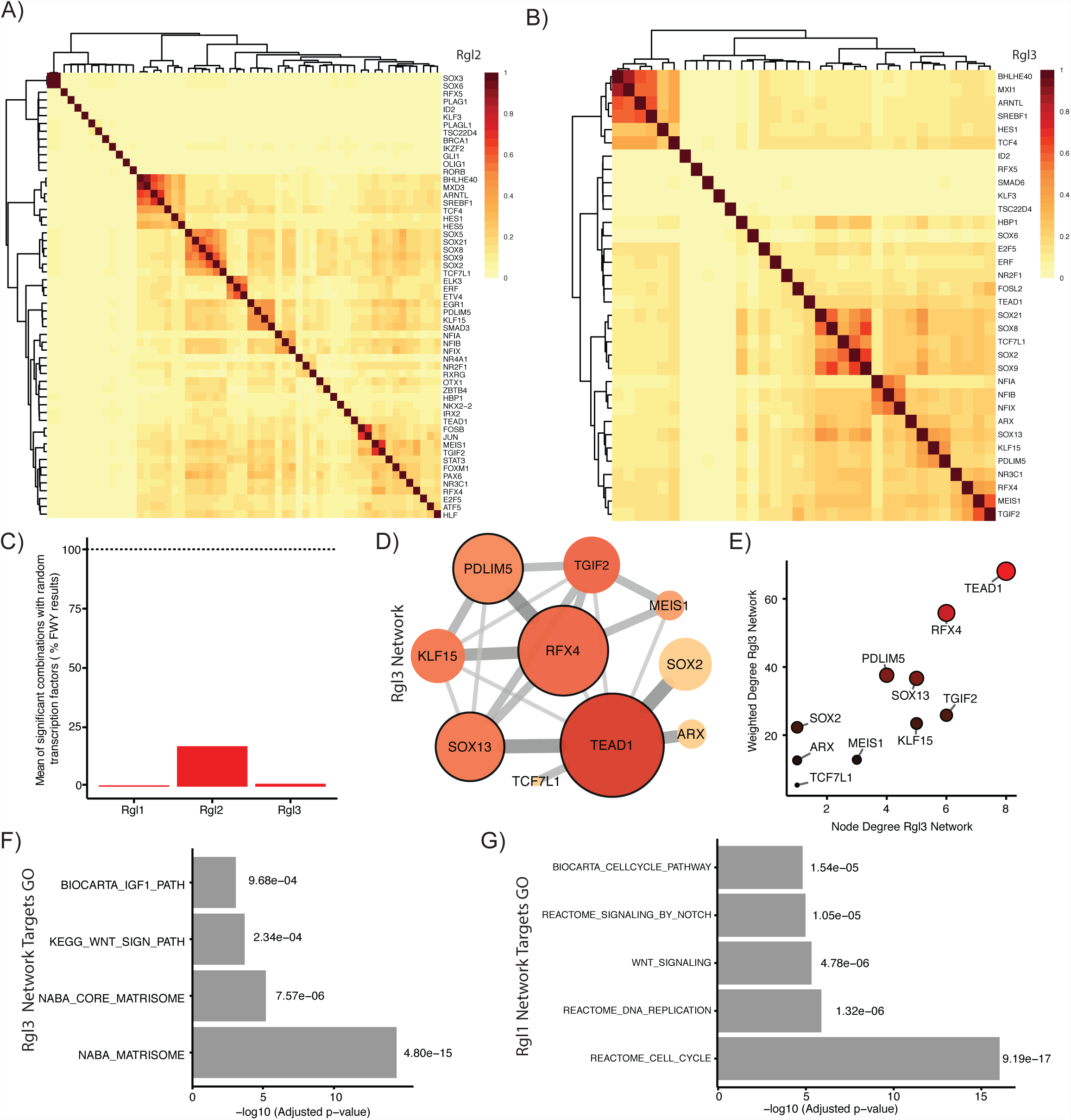
Combinatorial enrichment of transcription factors on Rgl2 and Rgl3. **(A)** Clustering of transcription factors expressed in Rgl2 by Jaccard index of shared target genes. **(B)** Clustering of transcription factors expressed in Rgl3 by Jaccard index of shared target genes. **(C)** Percentage of FWY significant combinations obtained with random selection of transcription factors compared to FWY results of Rgl1-3 transcription profiles. **(D)** Network representation of FWY analysis of Rgl3. Node color and size are proportional to node degree. Nodes with higher weighted degrees (core nodes) have a black border. Color intensity and width of the lines connecting the nodes are proportional to the interaction score for each transcription factor pair. **(E)** Plot comparing node degree and weighted node degree of the network obtained by FWY analysis of Rgl3. Color intensity and size are proportional to the weighted degree. **(F)** Gene enrichment of target genes for core transcription factors in Rgl3. Analysis was performed with MSigDB gene set C2 canonical pathways v5.0. **(G)** Gene enrichment of target genes for core transcription factors in Rgl1. Analysis was done with MSigDB gene set C2 canonical pathways v5.0.

**Figure EV7.**
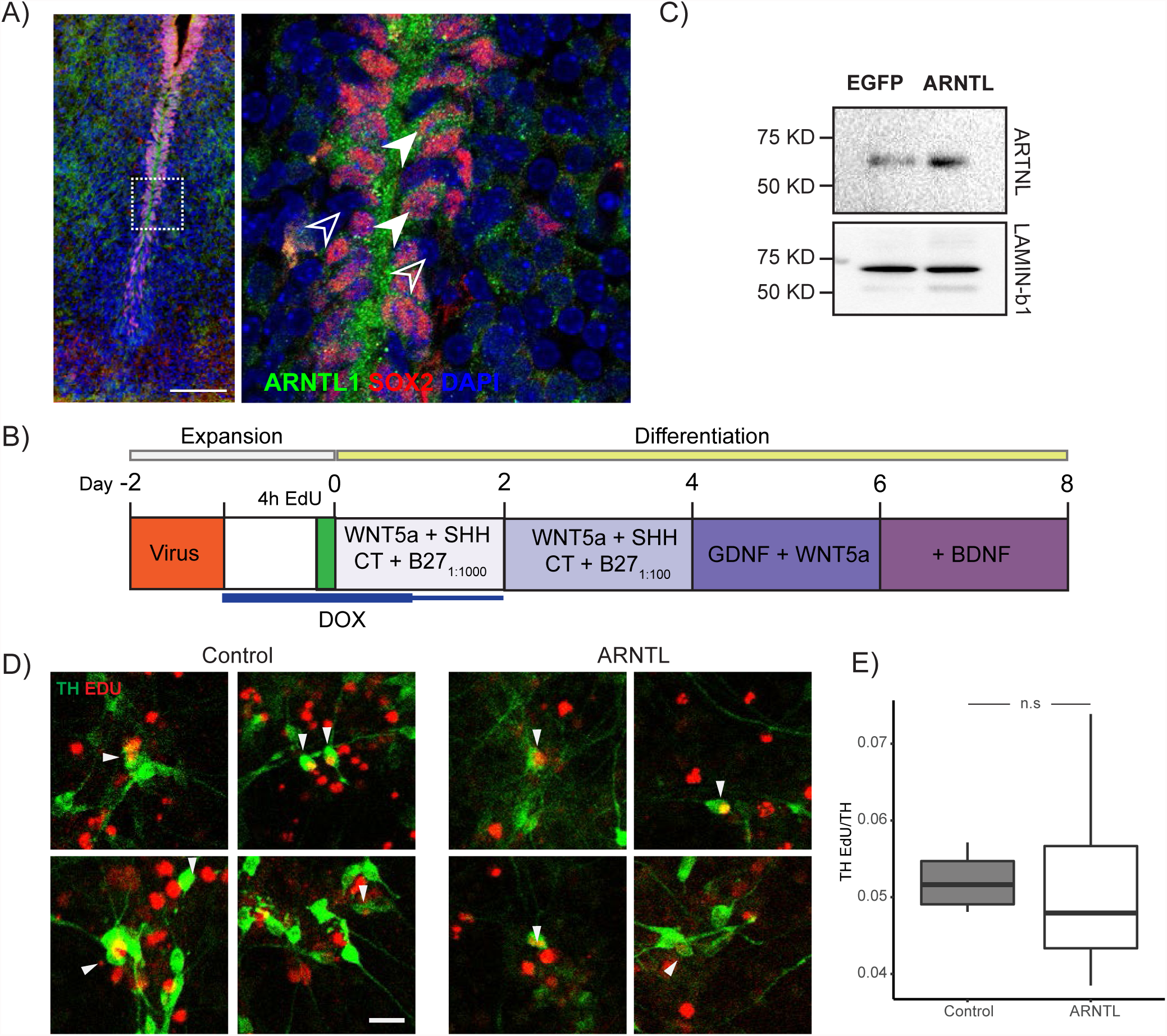
Overexpression of ARNTL in human neuroepithelial stem cells during dopaminergic differentiation. **(A)** ARNTL protein (green) is present in SOX2+ cells (red) in the developing mouse VM at E13.5. DAPI in blue. Scale bar, 100 μm. Image to the right magnified from box to the left. Full arrow heads, double positive SOX2 and ARNTL cells. Empty arrowheads, SOX2 and ARNTL negative cells. **(B)** Protocol for the overexpression of ARNTL in hLT-NES cells. **(C)** Increased levels of ARNTL were detected 24h after doxycycline treatment to LT-NES cells infected with ARNTL-lentivirus, compared to EGFP as control hLT-NES cells. LAMIN-b1 was used as loading control. **(D)** Representative immunofluorescence images corresponding to control and ARNT transduced hLT-NES cells at day 8. Cells having undergone mDA neurogenesis (arrowhead) are identified by the incorporation of EdU (red) in TH+ cells (green). Scale bar, 50 μm. **(E)** Quantification of mDA neurogenesis: TH^+^ and EdU^+^ cells relative to the total TH^+^ cells (n.s, non-significant, N = 4).

